# Breakage fusion bridge cycles drive high oncogene copy number, but not intratumoral genetic heterogeneity or rapid cancer genome change

**DOI:** 10.1101/2023.12.12.571349

**Authors:** Siavash Raeisi Dehkordi, Ivy Tsz-Lo Wong, Jing Ni, Jens Luebeck, Kaiyuan Zhu, Gino Prasad, Lena Krockenberger, Guanghui Xu, Biswanath Chowdhury, Utkrisht Rajkumar, Ann Caplin, Daniel Muliaditan, Ceyda Coruh, Qiushi Jin, Kristen Turner, Shu Xian Teo, Andy Wing Chun Pang, Ludmil B. Alexandrov, Christelle En Lin Chua, Frank B. Furnari, Thomas G. Paulson, Julie A. Law, Howard Y. Chang, Feng Yue, Ramanuj DasGupta, Jean Zhao, Paul S. Mischel, Vineet Bafna

**Author notes:** These authors contributed equally (Siavash Raeisi Dehkordi, Ivy Tsz-Lo Wong, Jing Ni).

## Abstract

Oncogene amplification is a major driver of cancer pathogenesis. Breakage fusion bridge (BFB) cycles, like extrachromosomal DNA (ecDNA), can lead to high copy numbers of oncogenes, but their impact on intratumoral heterogeneity, treatment response, and patient survival are not well understood due to difficulty in detecting them by DNA sequencing. We describe a novel algorithm that detects and reconstructs BFB amplifications using optical genome maps (OGMs), called OM2BFB. OM2BFB showed high precision (>93%) and recall (92%) in detecting BFB amplifications in cancer cell lines, PDX models and primary tumors. OM-based comparisons demonstrated that short-read BFB detection using our AmpliconSuite (AS) toolkit also achieved high precision, albeit with reduced sensitivity. We detected 371 BFB events using whole genome sequences from 2,557 primary tumors and cancer lines. BFB amplifications were preferentially found in cervical, head and neck, lung, and esophageal cancers, but rarely in brain cancers. BFB amplified genes show lower variance of gene expression, with fewer options for regulatory rewiring relative to ecDNA amplified genes. BFB positive (BFB (+)) tumors showed reduced heterogeneity of amplicon structures, and delayed onset of resistance, relative to ecDNA(+) tumors. EcDNA and BFB amplifications represent contrasting mechanisms to increase the copy numbers of oncogene with markedly different characteristics that suggest different routes for intervention.

## INTRODUCTION

Somatic copy number amplification (SCNA) of tumor promoting oncogenes is a major driver of cancer pathogenesis^1^. High copy amplifications (CN >8) are typically localized to specific genomic regions as focal amplifications fCNA)^2,3^. Currently, two main mechanisms for high copy number oncogene amplification predominate: extrachromosomal DNA (ecDNA)^4,5^ and breakage fusion bridge (BFB) cycles^6–8^. The independent replication of ecDNAs, their random segregation into daughter cells, and the positive selection for a higher (or appropriate) number of proliferative elements (e.g., oncogenes) provides the genetic basis for the rapid modulation of oncogene copy numbers in cells and explains much of the focal oncogene amplifications observed in cancer^4,9^. In addition, the circular shape of ecDNAs alter their chromatin accessibility, remodels their physical structure, and interactions of the DNA and its topological domains^10–13^. The remodeled structure changes the epigenetic landscape, generates new long-range cis-regulatory interactions, and enhances oncogene transcription to drive tumor pathogenesis. The rapid modulation of DNA copy number mediates resistance to targeted therapy via multiple mechanisms. These include: an increase in the number of copies through selection for higher numbers of ecDNA per cell ^14^, a decrease in copy number by reintegration of ecDNA into non-native chromosomal locations, or through the biogenesis of entirely new, compensating ecDNA^15^. Patients whose tumor genomes carry ecDNA are known to have worse outcomes relative to those with ecDNA(-) samples^5^. For these reasons, identification of ecDNA, and vulnerabilities specific to ecDNA(+) tumors remains an important challenge for understanding cancer biology.

The other main mechanism linked to high copy number oncogene amplification in cancer, Breakage-Fusion-Bridge (BFB) cycles, was proposed nearly 80 years ago by Barbara McClintock to explain patterns of genomic variation in irradiated maize cells ^16,17^. BFB cycles start during a bridge formation (usually between sister chromatids) as a stabilizing repair intermediate for DNA breaks or telomere loss (**Supplementary Figure 1**). Unequal mitotic separation of the bridged chromosomes creates an inverted duplication on one chromosome, and a deletion on the other. The broken ends result in continued BFB cycles until the telomere is re-capped. As with ecDNA, the lengthened chromosome may contain an oncogene that provides a proliferative advantage and selection for high copy number chromosomes creates rapid copy number amplification through successive BFB cycles. Other rearrangements might accompany BFB, confounding detection.

BFB cycles are an important driver of oncogene amplification in cancer. BFB-associated lesions were found in 41 of 45 tumor samples but rarely in normal fibroblasts^18^. HER2 positive breast cancers revealed a significant enrichment of BFB signatures and ecDNA within amplified HER2 genomic segments^19,20^. Experimental work has also revealed mechanistic aspects of BFB cycles. Mice that were deficient in both non-homologous end joining (NHEJ) DNA repair protein(s) and TP53 developed lymphomas that harbored BFB amplification of IgH/c-myc^21^. Anaphase bridge breakage was attributed to mechanical tension on structurally fragile sites, and the bridges were most frequently severed in their middle irrespective of their lengths^22^. Repeat=mediated genomic architecture surrounding the ERBB2 (HER2) locus was implicated in promoting BFB cycles^19^. Recently, lagging chromosomes were shown to initiate anaphase bridges, along with template switching at BFB junctions, and the bridge formation and breakage could lead to other catastrophic rearrangements^23^.

These results are indicative of a unique and important role for BFB cycles in cancer progression. However, the scope and extent of BFB amplification in cancer is not completely known, including the role of fragile regions in mediating breaks of the anaphase bridge and the consequent clustering of BFB structures. Moreover, it is not clear if the different modes of oncogene amplification including BFB cycles, ecDNA accumulation, and other intrachromosomal rearrangements are functionally interchangeable for cancer progression, if they amplify the same oncogenes, and whether they are prevalent in the same cancer subtypes. These questions require a systematic survey of BFB amplifications across many cancers, which in turn demands methods for reliable detection of BFB amplifications from genomic data.

Here, we addressed several of the aforementioned questions. Specifically, we developed a method, OM2BFB, for the detection of BFB and the characterization of their structure using BioNano Optical Genome Maps (OGMs). We validated the OGM based detection of BFB(+) amplification mechanisms through extensive cytogenetics experiments. Next, using OM2BFB as the standard, we benchmarked short-read based BFB detection, using our previously developed AmpliconSuite (AS) toolkit ^24,25^. We applied OM2BFB and AS on 1,538 whole genome samples to identify the location, scope, gene content and structural aspects of BFB based focal amplification. Finally, we integrated functional data including chromatin conformation, response to targeted drug treatment, gene expression, and patient outcomes to gain insight into how BFB cycles contribute to diverse cancer phenotypes.

## RESULTS

The pure breakage fusion bridge cycle can be described formally with a chromosomal loss, followed by a cycle of repeated anaphase bridge formations between (sister) chromatids, and bridge breaks during cytokinesis. See **Supplementary Figure 1** and **Methods** for a formal description. Sampling genomic sequences from a BFB-rearranged genome, and mapping them back to the human reference leads to a characteristic BFB signature of ladder-like copy number amplifications and an abundance of fold-back structural variations. These signatures are, however, not definitive, and many non-BFB amplifications, including ecDNAs, can also carry them. For example, the signature 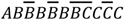, where each letter denotes a genomic segment, and the bars indicate inversions, shows a BFB signature, but is not BFB.

The OM2BFB algorithm utilizes whole genome optical genome map data, pre-processed by Bionano solve, to better identify foldback reads and copy number profiles. It utilizes the pre-processed data to locate focal amplifications carrying the BFB signature as candidate regions (**Figure 1A**^26^). For each candidate region, it then enumerates multiple possible BFB architectures, modifying methods that were previously developed, including by us, for short read whole genome sequences ^27–29^. Finally, OM2BFB scores each candidate architecture using a novel algorithm to estimate the (negative log-) likelihood of BFB cycle formation (**Methods**). High likelihood (low scoring) reconstructions are output, along with the score, in a stylized format (**Figure 1B**).

**Figure 1:**
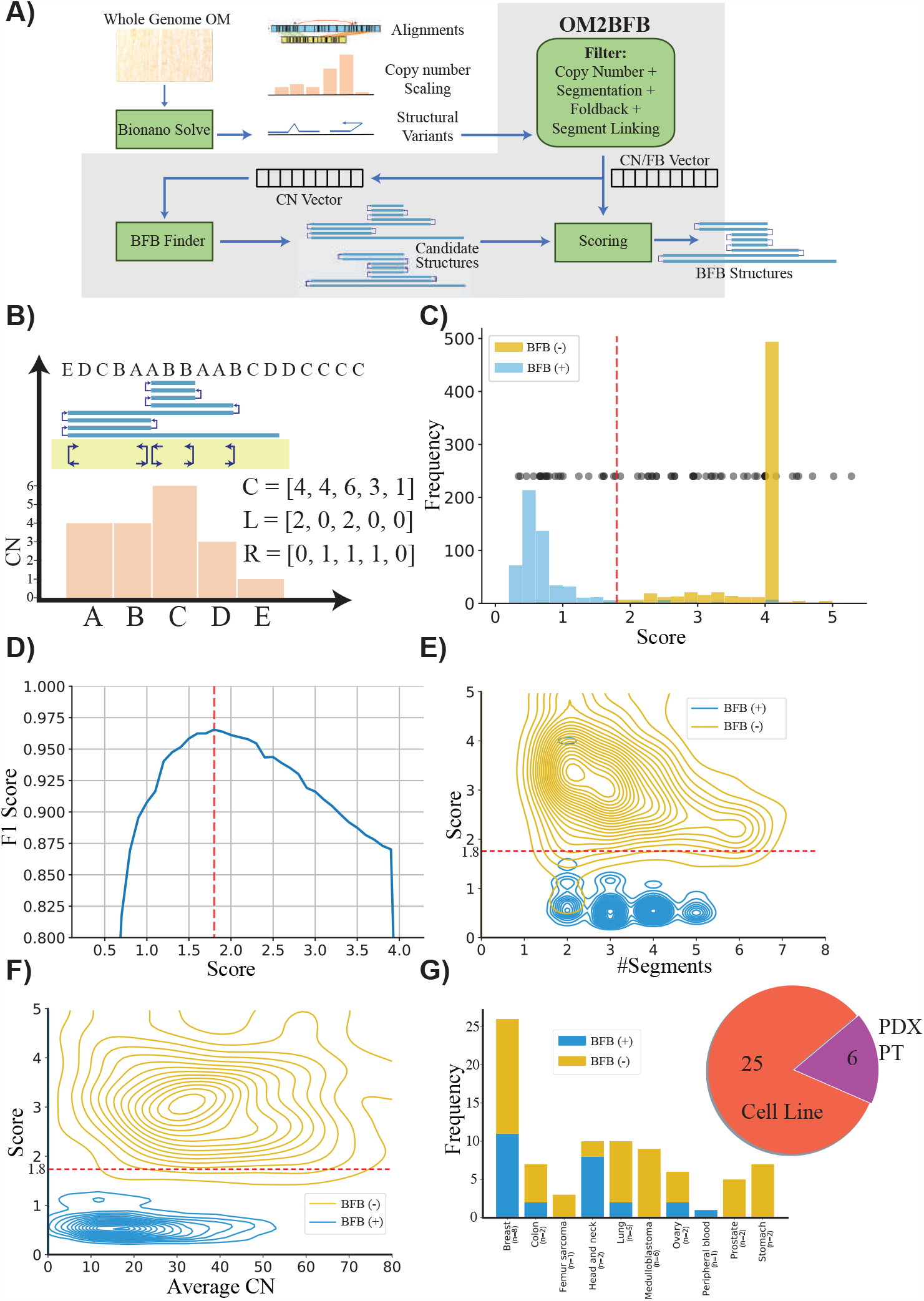
Detecting and reconstructing BFB amplicons using Optical Genome Maps. A).Workflow of OM2BFB pipeline depicting the sequential steps involved in the analysis (Methods). B).Schematic representation of OM2BFB output. The axes display genomic coordinates (x-axis) and copy-number (CN) (y-axis), with a separate track showing fold-backs. Orange bars represent segments with their respective copy numbers, navy arrows on the yellow background indicate foldback reads, and the blue rectangle on top represents the BFB structure described by an ordering of segment labels. The proposed BFB structure can be read by marking the segments traversed by the blue line, starting from the centromeric end. The vectors, C, L, and R refer to the copy number, left-, and right-foldback SVs that support the BFB. C).Distribution of OM2BFB scores for 681 simulated BFB negative (BFB(-)) cases and 595 simulated positive (BFB(+)) cases. Notably, 490 negative cases did not meet the filtering threshold and were reported with a high score (4) by OM2BFB. Black dots (placed at an arbitrary position on the y-axis for ease of visualization) represent the OM2BFB scores of 84 amplicons obtained from 31 cell-lines. The red line at 1.8 marks the threshold from (D) for defining BFB(+) from BFB(-) cases. D).The F1 scores of OM2BFB, measured for different score cut-offs. The highest F1 score was achieved at a threshold of 1.8, and that score was selected as the threshold for separating BFB (+) from BFB(-) cases. E).Distribution of OM2BFB scores across the number of segments in the simulated cases. The BFB(+) sample scores are independent of the number of segments, while BFB(-) samples reveal a slight bias of decreasing scores with higher number of segments. F).Distribution of OM2BFB scores across the average segments’ copy numbers in the simulated cases shows independence between the score and average copy number. G).(Right) Distribution of OGM data types from 31 samples that include cancer cell lines, PDX models, and primary tumors (PT). (Left) Distribution of BFB(+) and BFB(-) cases across different cancer subtypes.

We tested the performance of OM2BFB using extensive simulations. To enable this, we developed a BFB simulation method and used it to simulate 595 BFB-positive data sets, spanning a large number of segments and copy numbers. Additionally, we used ecSimulator^30^ to generate 681 BFB-negative cases, including ecDNAs. The positive and negative examples largely overlapped in the parameter space (**Supplementary Figure 2**). However, ecDNA based amplifications had higher copy numbers relative to BFB amplifications, consistent with current knowledge. OM2BFB scoring clearly separated the positive and negative examples (**Figure 1C**). The maximum F1 score, describing the harmonic mean of the precision and recall, equaled 0.96 at a score-cutoff of 1.8 (**Figure 1D**). Notably, 490 of the BFB negative examples were trivial in that they either failed the initial filtering or represented a single segment with left and right fold-backs (deemed to be an ecDNA; **Figure 1C**). Nevertheless, a significant number of the negative examples passed the initial filters and some would have been identified as BFB by available BFB calling tools using currently accepted metrics. We noted that the score distribution stayed consistent with changes in the number of segments from 1 to 7, and also with variation in the average copy number from 5 to 80 (**Figure 1E**,**F**). Different components of the score, corresponding to segmentation, and fitting of the observed copy numbers and foldbacks to the candidate structures all contributed to the final performance (**Supplementary Figure 3**). The small numbers of false negative examples could be attributed to missing fold-back reads, while false positives were mainly due to ecDNA structures that strongly resembled BFBs in their shape (**Supplementary Figure 4**)

We next applied OM2BFB to 84 amplicons using OGM data from 31 samples, including cancer cell lines, patient-derived xenograft (PDX) models, and primary tumors (**Figure 1G** and **Supplementary Table 1**). Using the cut-off score of 1.8, 61 cases were identified as being BFB(-), and 23 as BFB(+), with high numbers of positive occurrences in Her2+ Breast cancer and Head and Neck cancer lines.

Unlike for ecDNAs, fluorescence in situ hybridization (FISH) is not a fool-proof method for detecting BFB structures, even during metaphase, when the chromosomes can be delineated. The presence of a homogeneously staining region (HSR) on a native chromosomal location is characteristic of BFB(+) structures, but every native HSR may not be amplified via BFB. Conversely, HSR amplification on a non-native chromosome with no amplification on the native chromosome is indicative of a BFB(-) structure, for example, through ecDNA formation and reintegration at a non-native locus. A BFB(+) structure may also occur when a HSR amplification is present on both a non-native chromosome and a native chromosome, indicating that the BFB structure broke away and translocated to another chromosome. We cytogenetically visualized 27 (19 metaphase, 8 interphase) structures that had previously been scored by OM2BFB, using a probe from the amplified region (**Supplementary Table 1**). In metaphase cells, we used the criteria of “HSR amplification on a native chromosome” and appearance of left-foldbacks and right-foldbacks in the optical genome mapping (OGM) data to validate 7 BFB(+) and 12 BFB(-) predictions. 6 of the 7 BFB(+) structures (**Figure 2A**,**B** and **Supplementary Figure 5A-E**) and 12 of 12 BFB(-) structures were predicted accurately by OM2BFB (**Figure 2C**,**D** and **Supplementary Figure 5F-K**). The BFB(+) structures included simple cases, such as a *MYCL1* amplification in *THP1* (**Figure 2A**), but also complex ones, including a 10+Mbp BFB amplification in the HARA cell line that amplified *PDHX1* on chr11p (**Figure 2B**). The OGM data detected additional translocations from the short arm (p arm) to the long arm (q arm) of chromosome 11, and chr11q contained an amplification of *CCND1*. Using multi-FISH probes for *PDHX1* (red), *CCND1* (green), and chr11 centromere (yellow) (**Figure 2C**), we confirmed that *PDHX1* was amplified as a BFB on the native locus but also translocated to the q arm where it co-amplified *CCND1*.

**Figure 2:**
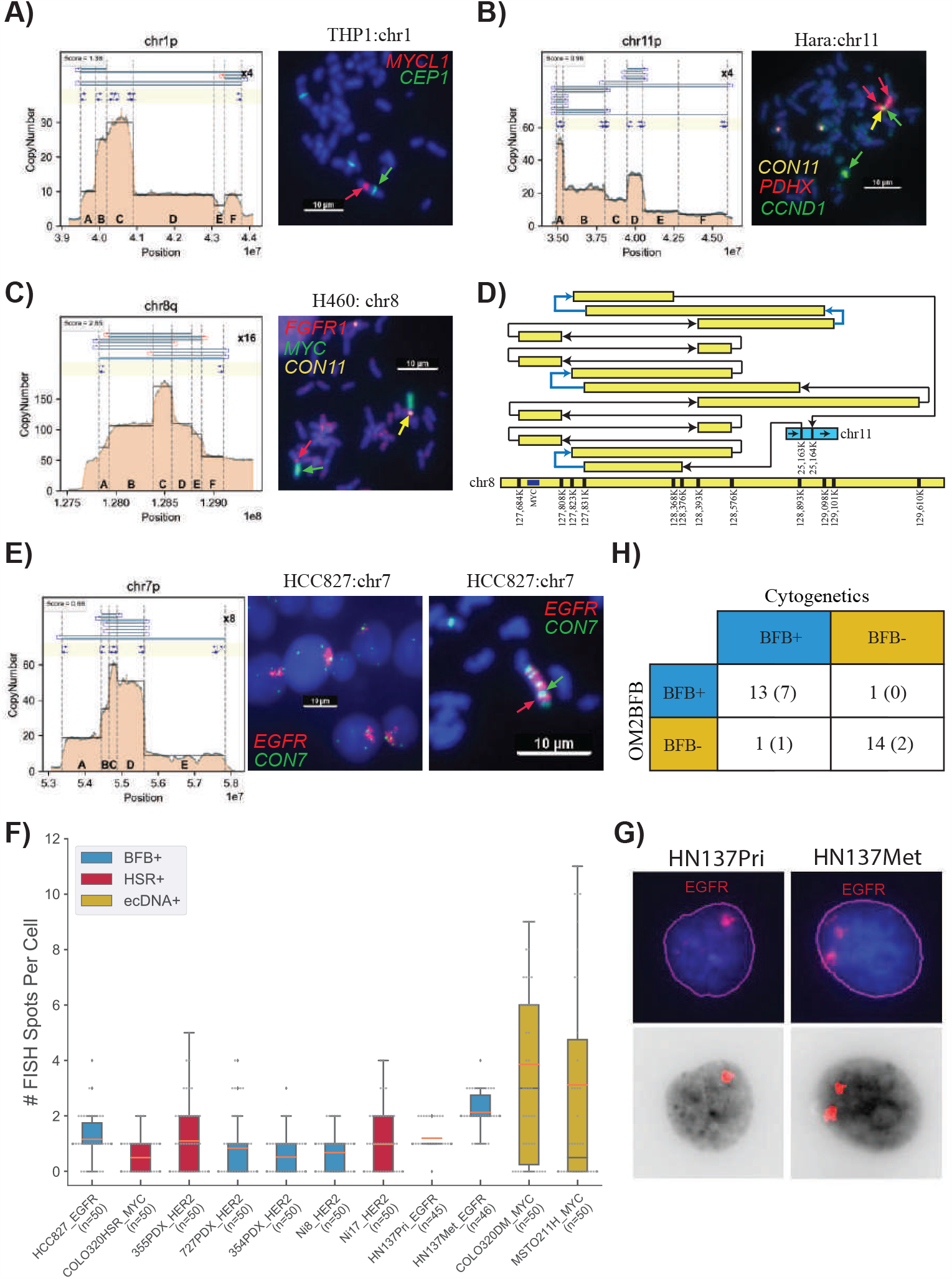
A).Metaphase FISH image for the cancer cell line THP1, showcasing amplification on a native chromosome (chr1) with a score below 1.8. CEN1 is a probe for the centromere on chr1. B).Metaphase FISH image for the cancer cell line HARA, displaying amplification on a native chromosome arm (chr11p) and also co-occurrence of HARA and CCND1 on the q-arm, suggesting a duplicated translocation of the BFB site from the p arm to the q arm. C).Metaphase FISH image for the cancer cell line H460, demonstrating amplification on both native (chr 8) and non-native (chr 11) chromosomes with scores exceeding 1.8. D).Visualization of HSR (homogeneously staining region) amplification on chr 8 (MYC) integrated in chr 11 in the H460 cell line. Blue arrows represent foldback reads within the structure. E).Metaphase and Interphase FISH images for the cancer cell line HCC827, exhibiting amplification on a native chromosome with a score below 1.8. F).Distribution of FISH Foci Count among cases with Interphase FISH images, highlighting lower number of foci and also lower variance in the number of foci in BFB and HSR cases compared to ecDNA cases. G).Visualization of EGFR foci in interphase cells from HN137Pri and HN137Met lines. The top panel shows the original FISH image, while the bottom panel shows the computationally detected foci. H).Summary of cytogenetic validation of OM2BFB calls. The number in parentheses refers to the number of interphase samples.

OM2BFB gave a high score (>1.8) for all cytogenetically negative cases. For example, it gave a score of 2.93 to the *FGFR2* amplicon in SNU16 (**Supplementary Figure 5K**). The amplicon is an ecDNA with extensive foldback reads. A more interesting example was a Myc amplicon in the cell line H460. Despite extensive fold-backs, the OM2BFB score was 2.65 (**Supplementary Figure 5G**). We performed a multi-FISH experiment probing for Myc (green), FGFR1 (red; near chr 8 centromere), and cen11 (yellow).Cytogenetics analysis clearly indicated two HSRs, one at the native locus on chr8, and the other on chr11 (**Figure 2C** and **Supplementary Figure 5G**). A careful reconstruction revealed a complex rearrangement of chr8 genomic segments with an insertion in chr11 (**Figure 2D**). The amplicon contained multiple foldback SVs, but the patterns did not support a BFB origin. This likely represents a case of an ecDNA that re-integrated back to the native locus, but also to a nonnative locus. The result reaffirms that amplicons with high copy numbers and foldbacks could be non-BFB related HSRs.

The *MCL1* amplicon in chromosome 1 of the COLO320 line had previously been reported to carry inverted fold-back reads^31^. FISH imaging suggested an HSR on chr1q, but also found the long arm to be detached from chr1p (**Supplementary Figure 5H**). Amplification was also observed on non-native chromosomes. The OM2BFB reconstruction shows a simple structure that is missing foldbacks required to complete the BFB cycle (**Supplementary Figure 5H**), suggesting that this is a BFB(-) case, but one that arose with anaphase bridge formation and breakage.

### BFB structures show lack of heterogeneity in interphase cells from patient tumors

We tested OM2BFB in two separate patient data sets: the first dataset was a cohort of 6 samples, including primary tumor and patient-derived xenografts from individuals who had HER2(+) breast cancer with brain metastases (BCBM) ^32,33^. The second data set consisted of untreated primary (HN137-Pri) and metastatic (HN137-Met) cell lines derived post-surgery from a patient with metastatic head and neck (HN) cancer^34^. Optical genome map data was acquired for these samples and analyzed using OM2BFB.

OM2BFB predicted 7 BFB amplicons in the BCBM data, including 2 containing HER2 (**Supplementary Table 1 and Supplementary Figure 6**). In the HN137 data from one patient, OM2BFB identified 8 BFB amplicons. EGFR was amplified at a BFB site in both HN137Pri and HN137Met cell lines but increased in copy number from CN=10-12 (Pri) to CN=20 (Met) (**Supplementary Figure 7**). Moreover, a BFB containing *YAP1* and *BIRC2* with high (CN=40) amplification was present in HN137Met, but absent in HN137Pri. A BFB on chr11q, containing *CCND1*, was observed in both Pri and Met. A chr3q BFB containing *EPHA1* was observed in HN137Pri, but not in Met. Finally, the copy number of a chr18p BFB was reduced from 30 to 10 and reduced in span.

As metaphase FISH was not possible for patient tumor cells, we used interphase analysis. We developed methods to segment the nuclei, identify and count FISH signals per nucleus. For BFB cycles and HSRs, we would expect to see lower counts *and* lower variability in the number of foci from cell to cell, in contrast to the higher counts and variance between cells for ecDNA(+) foci. We confirmed that the interphase foci had this property for known BFB in HCC827 (EGFR), HSR in Colo320HSR (Myc), ecDNA in Colo320DM(Myc) and MSTO211H (Myc) (**Figure 2E**,**F)**. Utilizing this approach, OM2BFB predictions were consistent with interphase FISH in 7 of 8 BCBM BFBs, with one false negative prediction, and in 2 of 2 HN137 cell lines with *EGFR* amplifications (**Figure 2F, Supplementary Figure 6** and **Supplementary Table 1**). Intriguingly, DNA FISH probes for *EGFR* showed two stable foci per cell in HN137Met in contrast to a single HSR in HN137Pri (**Figure 2F**,**G**), explaining the doubled copy number in the metastatic line. The homogeneity of BFB structure between HN137-Pri and HN137-Met lines (**Supplementary Figure 7**) suggested a stable translocation from primary to metastatic cells. To explore this further, we additionally analyzed the HN137 cell-lines in metaphase, and confirmed the change in the number of chromosomal foci from one to two. Remarkably, a small subset of HN137Met cells (5 of 60) displayed EGFR on ecDNA (**Supplementary Figure 8)**, raising the possibility that the translocation is mediated by ecDNA.

Taken together, the cytogenetics revealed that the presence of focal-amplification and foldback reads are not, by themselves, sufficient to predict BFB. They confirmed the utility of OM2BFB as a sensitive (13/14=92.8%) and precise (14/15=93%) method for BFB prediction (**Figure 2H)**. They revealed that BFB driven amplicons are followed by additional rearrangements, including translocation to other loci and ecDNA formation. This is consistent with earlier results that suggest that anaphase bridge formation and breakage can be a precursor to gross DNA instability^23^, including ecDNA formation^35^. At the same time, there are many cases where BFB cycles result in stable focal amplification on the native chromosome, with low cell to cell heterogeneity.

### BFB amplifications are ubiquitous in multiple cancer subtypes

The Amplicon Suite (AS) pipeline, comprising Amplicon Architect and Classifier, has been validated for ecDNA detection, but not for BFB detection, due to a lack of positive and negative examples. We compared the AC results on the same 83 amplicons initially scored by OM2BFB (**Figure 1C**). Of the 23 BFB predictions made by OM2BFB, AC predicted 14 (**Supplementary Table 1**). It predicted some BFB(+) amplifications as BFB(-), mainly due to a lack of foldback read prediction using short-reads, especially when the coordinates between the fold-back reads spanned many Kb (**Supplementary Figure 9**). On occasion, the persistent reuse of left and right foldback reads suggested a circularization of the BFB structure into an ecDNA. Importantly, however, AC did not make a single BFB call in the 61 amplicons that OM2BFB labeled as being BFB(-) by OM2BFB (**Supplementary Table 1**). Because the AC BFB calls showed high precision (few false positives), we utilized it to understand population distribution of BFB structures in larger cancer genome repositories.

We executed Amplicon Architect (AA) followed by AmpliconClassifier (AC) on 1,538 genomes from The Cancer Genome Atlas (TCGA), on 305 genomes consisting of premalignant Barrett’s esophagus and esophageal cancer samples (BE)^36^, and on 270 samples from the Cancer Cell Line Encyclopedia (CCLE)^37^ (**Figure 3A**). AC identified 371 BFB amplicons, including 258 in primary tumors, located across nearly every chromosome (**Figure 3B**). Notably, the incidence in TCGA was markedly lower than in the BE and CCLE data. This can be attributed to the higher incidence of BFB in esophageal and lung cancers, and the high proportion of lung samples in CCLE data (**Supplementary Table 2**).

**Figure 3:**
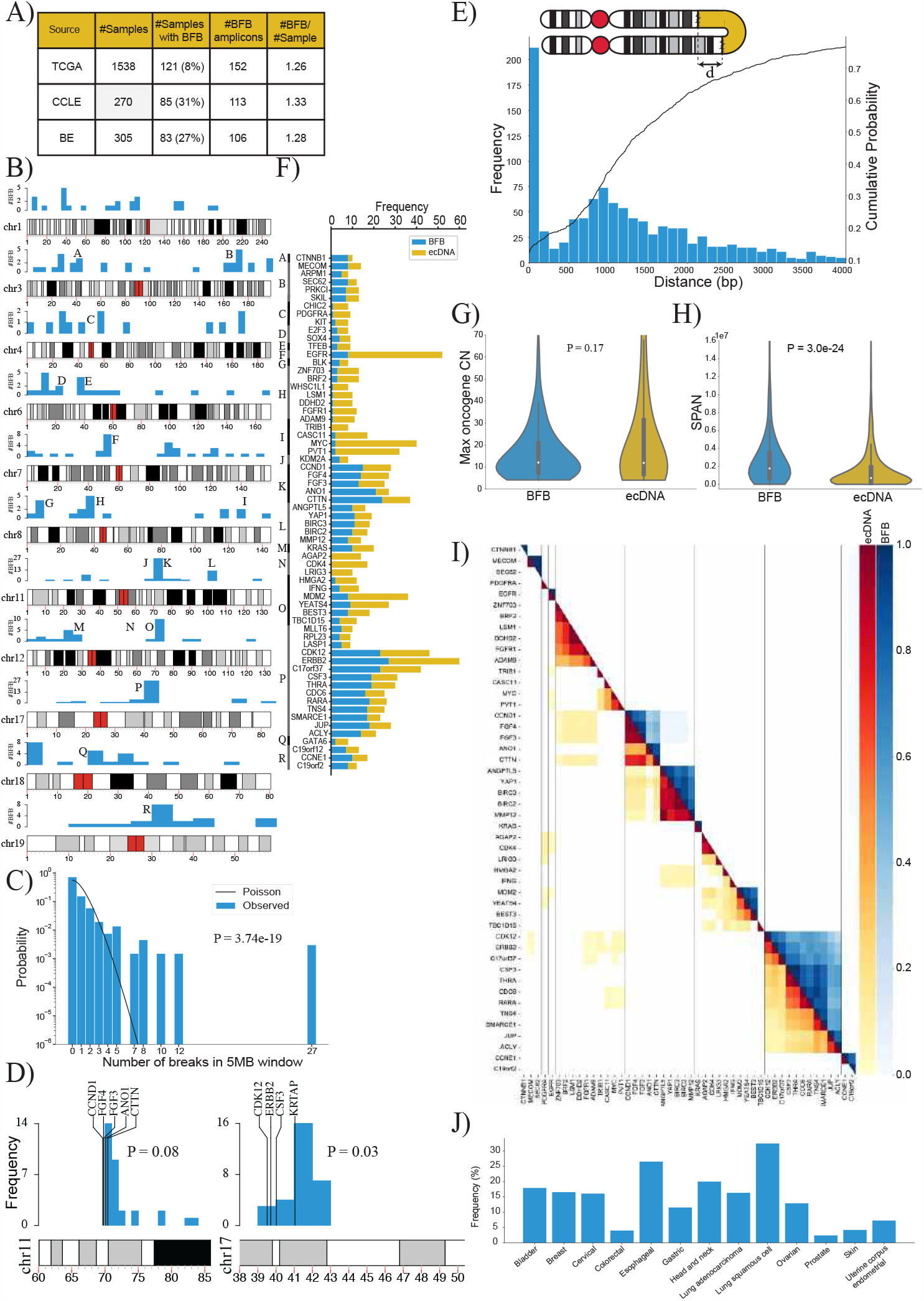
A).Summary of the number of BFB(+) samples among 2,113 whole genome samples tested for BFB amplification using AmpliconSuite. The data was collected from three data sets: TCGA, BE^36^ and CCLE^37^. B).Locations of the first (most telomeric) break of the 371 BFBs in the human genome (hg38). Chromosomes with fewer than 12 BFBs are not shown. C).The distribution of BFB occurrences (most telomeric break) in 5 Mbp windows compared against the Poisson distribution to test for randomness (p-value: 3.7e-19, KS test). D).The randomness of the first break in a 10Mb region, telomeric to an amplified oncogene. Left panel: 29 BFBs on chr11 containing CCND1; Right panel: 30 BFBs on chr17 (ERBB2). E).Distribution and cumulative distribution of the distance (d) between fold-back reads. F).Frequencies of the mode of amplification (BFB versus ecDNA) in oncogenes that are amplified at least 8 times in all datasets combined. G).Violin plot showing the distributions of the maximum oncogene copy number between BFB and ecDNA amplicons (p-value = 0.17 with Ranksum test). H).Violin plot showing the distributions of amplicon length (SPAN) between BFB and ecDNA amplicons (p-value = 3.0e-24 with Ranksum test). I).Co-occurrence patterns of amplified oncogenes. Color-coded entry for (i,j) measures the fraction of times genes i and j were both amplified when either gene was amplified. The lower triangle shows ecDNA co-occurrence patterns and the upper triangle shows BFB co-occurrence patterns. J). Distribution of BFB amplicons over different cancer subtypes. BFB amplicons were not found in brain and CNS related cancers, but were most abundant in lung and head and neck cancers.

### BFB locations are highly dispersed, but not random

BFB structures were identified in multiple chromosomes, and did not show any preference for centromeric or telomeric locations (**Figure 3B**). We partitioned the genome into 5Mb bins and marked the ones that carried BFB amplifications. Intriguingly, the BFB locations were not distributed randomly in those binds (**Figure 3C**; p-value 3.7e-19, KS test). For example, a bin comprising region K on chromosome 11 containing the genes *CTTN* and *CCND1* carried 27 of 371 BFB amplicons. Furthermore, while chromosomes 5 and 13 had fewer than 7 BFB events, chromosomes 3, 7, and 11 had more than 30 BFB events each. Multiple hypotheses explain the non-random distribution. The initiating break could occur at a fragile region of the genome. A recent result points to fragility in the *KRTAP1* region, 1.12Mb telomeric to *HER2* as a cause for recurrent BFB amplification of *HER2*^38^. A second hypothesis is that the initial break is random, but subsequent amplification of an oncogene and positive selection for higher copy numbers leads to recurrent BFBs in specific genomic regions. To test these two hypotheses, we chose 7 genomic regions that were recurrently amplified via BFB at least 7 times. For each such region, we looked at 10 1Mb windows telomeric to an oncogene to test if the breaks were preferentially clustered. In all cases but one, the breaks were randomly distributed, connected only by the fact of sharing a common amplified oncogene (**Supplementary Figure 10, Supplementary Table 3**). The window containing the *KRTAP1* gene was indeed preferentially selected for HER2+ BFBs (**Figure 3D**; 16/30 breaks, nominal p-value 0.03; see methods). The *CCND1*-*CTTN* gene cluster on chromosome 11 also showed preferential break in a window immediately downstream, but did not reach the p-value=0.05 level of significance. In the other 5 cases, there was no preference for any window in the 10Mb region. While our data is sparse, the results suggest that an initial random breakage followed by positive selection for oncogene amplification can lead to a non-random distribution of BFB locations. Similarly, while the anaphase bridge breaks due to mechanical tension, a break immediately downstream of the oncogene is preferentially selected.

We observed that the distance between the fold-backs reads has a long tail, with over 65% of fold-back distances greater than 1000bp (**Figure 3E**). Moreover, the distribution was possibly under-estimated, given the ascertainment bias. The result suggests that fusion-bridge formation requires proximity, but not palindromic sequence.

### BFBs amplify the same oncogenes as ecDNA but are structurally different

We investigated the oncogenes amplified by BFB across all sample types. Consistent with the ‘random breakage with selection’ hypothesis, the BFB cycles amplified a large number of oncogenes. Intriguingly, the oncogenes amplified via BFB strongly overlapped with oncogenes amplified by ecDNA. Of the 52 oncogenes amplified at least 4 times on BFB cycles, 49 were also amplified as ecDNA at least once (**Figure 3F**; **Supplementary Table 4**). In contrast, 19 (20%) of the 96 genes that were amplified at least 4 times as ecDNA were not found to be BFB amplified (p-value 0.03, Fisher Exact Test). These included *CDK4, MDM4, FGFR3*, and others. These results support the hypothesis that while BFB amplification is one of many precursor events that lead to ecDNA formation, ecDNA can be formed through BFB-independent events, including episome formation and chromothripsis ^9,23,35^.

Positive (*trans-)*selection for ecDNA can occur if they carry regulatory elements that serve as a ‘roving enhancers’ for genes on other ecDNA or chromosomes^39,40^. In contrast, the chromosome bound BFB structures, with locally mediated rearrangements, are unlikely to hijack distal enhancers. We hypothesized that selection for BFB structures was largely mediated by amplification of oncogenes or *cis*-regulatory elements. Consistent with this hypothesis, we observed that only 5% (21 of 371) BFB amplicons did not carry a known oncogene (**Supplementary Table 2**) while over 10% (80 of 759) of ecDNA amplicons were oncogene free (p-value 0.003; Fisher exact test). The maximum oncogene copy numbers were, however, similar for BFB and ecDNA structures (**Figure 3G**).

While BFB and ecDNA amplified the same genes, the amplicon structures were different. BFB amplifications had a larger span compared to ecDNA (**Figure 3H**; mean span 2.8Mbp versus 1.5Mbp; p-value 3.0E-24). Surprisingly, despite BFB structures having a large span, they did not co-amplify more proximal oncogenes than ecDNA. In fact, ecDNA structures were often multi-chromosomal, and co-amplified distant oncogenes (**Figure 3I**, lower triangle). Such co-amplification was rare in BFB (**Figure 3I**, upper triangle), because it would require the presence of independent BFB amplifications, or the formation of translocation bridges^41^.

Similar to ecDNA, BFB amplicons were found in multiple cancer subtypes (**Figure 3J**). Certain subtypes were more likely to carry BFB amplifications, including lung, esophageal, and head and neck cancers. However, BFBs were rare in brain cancers, where ecDNA amplifications are prevalent ^5,13^.

### BFB amplified genes show lower variance of gene expression, with fewer options for regulatory rewiring

BFB cycles generate stable chromosomal amplifications and are likely to function differently from the more mobile ecDNA (and even the ecDNA re-integrated as HSRs). Circularization and other rearrangements have been shown to change the topologically accessible domain (TAD) structure for ecDNA, changing their regulatory wiring, including the hijacking of distal enhancers^12,42^. In contrast, fold-backs are the primary source of rearrangements in BFB cycles. While foldbacks also change the conformation, they would be less likely to bring distal regions in close proximity. To test this hypothesis, we used HiChIP to measure chromatin interactions involving the enhancer- and promoter-associated mark H3K27ac on focal amplifications in the cell-line COLO320DM. The cell line amplified *MYC* (chr8) on ecDNA^43^. It carries another focal amplification on chr1, that originated with a duplication inversion characteristic of BFB^31^. Consistent with genomic rearrangements, the HiChIP interactions maps on both ecDNA and BFB regions showed a remarkable change in topological structure, when compared to the matching genomic regions in the control GM12878 line (**Figure 4A**,**B**). For example, the rightmost foldback in the Chr1 amplicon topologically separates the telomeric region from the BFB region in COLO320DM, but not in GM12878. Similarly, the extensive rearrangements in the chr8 ecDNA create distant interactions in COLO320DM that are absent in GM12878. We next identified significant chromatin interactions in the two structures using NeoLoopFinder^44^. The ‘neo-loops’, marked by black spots, are interactions mediated by genomic rearrangements in the cell line, while the blue spots (‘loops’) are interactions attributable to conformational change (**Figure 4A**,**B)**. In both amplicons, we identified a larger number of distal (off-diagonal) interactions relative to GM12878. The number of distal H3K27ac-region interactions in the BFB amplification was larger relative to GM12878, but significantly lower than for the ecDNA amplification (**Figure 4C**; p-value 5.95e-05, Peacock (2d-KS) test). Thus, BFB driven amplifications (unlike with ecDNA) could have fewer options for rewiring of the regulatory circuitry.

**Figure 4:**
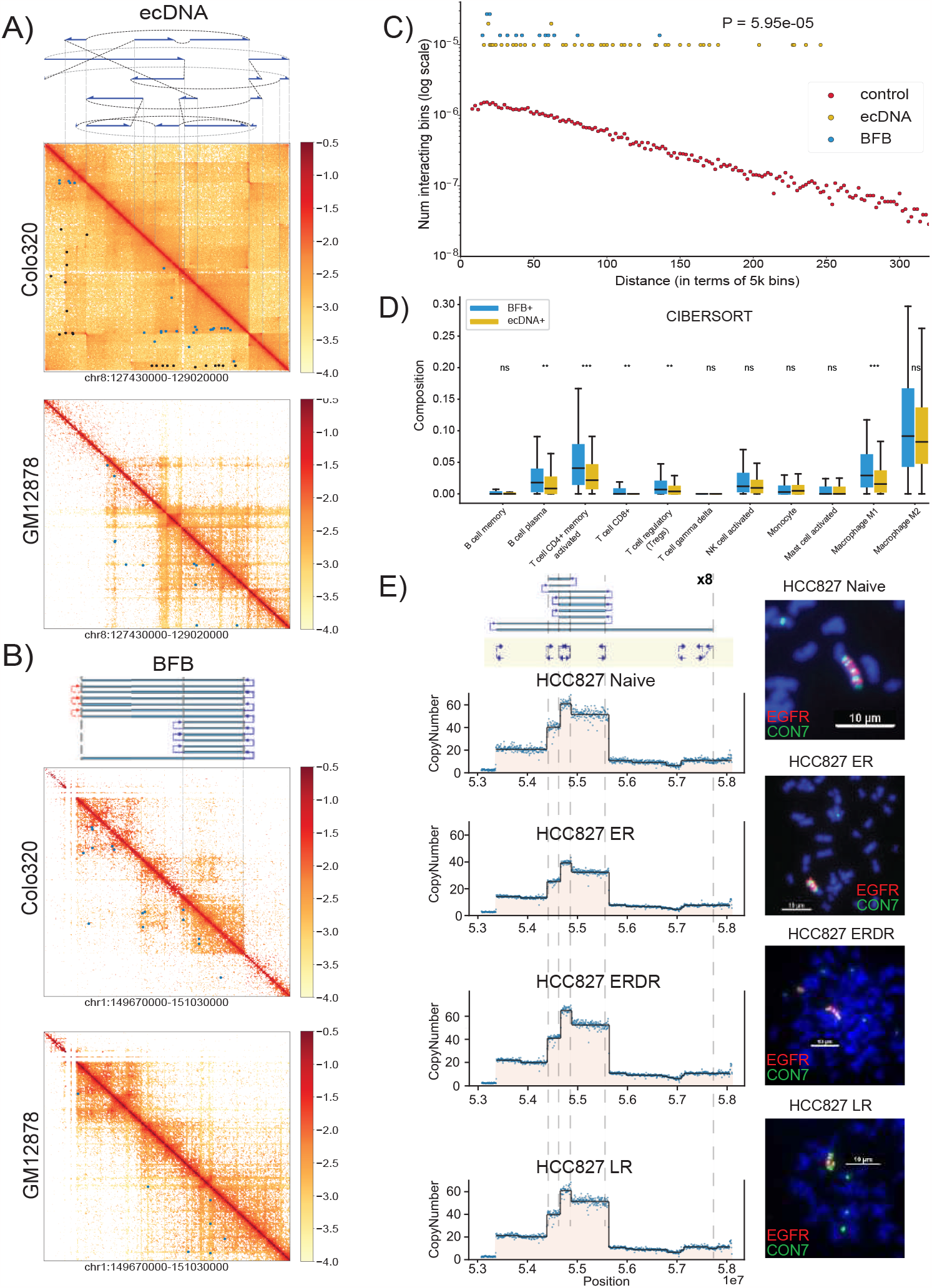
A).Top: The structure of the *MYC*-amplified ecDNA from the COLO320DM cell line. Blue arrows indicate genomic segments from chr 8 amplified on the ecDNA; black dashed lines indicate SV breakpoints directly connecting two remote genomic segments; gray dashed lines indicate templated insertions of segments involving other chromosomes. Middle: Normalized HiChIP contact map of COLO320DM at the ecDNA locus. Blue spots indicate significant chromatin interactions identified by NeoLoopFinder; while black spots indicate “neoloops” proximal to SV breakpoints and likely to be formed due to the genomic segments coming together in the cell line. Bottom: Normalized HiChIP contact map of GM12878 at the identical chr8 locus. Blue spots indicate significant chromatin interactions. B).Top: The inferred structure of the BFB-like focal amplification from the COLO320DM cell line. Middle: Normalized HiChIP contact map of COLO320DM at the BFB locus. Blue spots indicate significant chromatin interactions identified by NeoLoopFinder. Bottom: Normalized HiChIP contact map of GM12878 at the identical chr1 locus. Blue spots indicate significant chromatin interactions. C).Distribution of HiChIP interaction frequencies in ecDNA and BFB-driven amplifications. For a specific genomic distance d (x-axis), the dot represents the fraction, among all pairs of genomic windows separated by d, of pairs with significant HiChIP interactions. D).Differences in immune cell subtype compositions in BFB(+) cancers (n=76) versus ecDNA(+) cancers (n=297). (* indicates p<0.05; ** ⇒ p<0.01; *** ⇒ p<0.001). E).Targeted Therapy Resistance of the HCC827 Cell Line continuing EGFR amplified within a BFB event. The top panel shows the BFB architecture in the HCC827 naive cell line along with metaphase FISH images. Resistance formation to Erlotinib (ER) maintains the BFB amplicon structure, but the bulk copy number is highly reduced. The copy number and the proportion of cells carrying the BFB signal are restored after drug removal (ERDR). No changes were observed in the BFB amplification in Lapatinib drug resistant (LR) line.

We also explored the normalized gene expression data from the cancer genome atlas for genes amplified on BFB and ecDNA. The expression of BFB amplified genes increased with copy number, similar to ecDNA (**Supplementary Figure 11)**, but there were important differences in transcription of other genes, especially relating to the immune response. EcDNA amplification has previously been associated with lower immune activity^5,45^. Using previously estimated cell type composition based on transcript evidence^45^, we found that BFB(+) samples had increased concentration of multiple immune cell-types, including cytotoxic T cell (CD8+) levels and pro-inflammatory macrophages, relative to ecDNA(+) samples (**Figure 4D**). Remarkably, BFB(+) samples also showed increased expression of tumor checkpoint genes, including *BTLA, CTLA4, PD-L1/CD274* (**Supplementary Figure 12)**. Our results suggest that cancers with BFB amplification have lower immunosuppression than ecDNA amplifications, and might be more susceptible to checkpoint target engagement.

EcDNA(+) cells can rapidly modulate their copy number in response to changes in environment. For example, in a glioblastoma model, ecDNA integrated into chromosomes with reduced copy number upon drug treatment, and rapidly reappeared upon drug removal^43^ To test the mechanism of targeted therapy resistance in BFB-mediated amplifications, we obtained naive, drug resistant (Erlotinib and Lapatinib), and drug removed versions of the cell line HCC827, which amplifies *EGFR* on a BFB. Metaphase FISH confirmed the stable BFB amplicon in naive, drug resistant, and drug removed lines (**Figure 4E)**. Remarkably, the *EGFR* copy number in Eb resistant lines was 30% smaller than the naive and drug removed lines. These results were supported by DNA FISH experiments, which showed that the BFB was present in resistant cells, but only in 10 of 29 cells. In contrast, the amplification was universally observed in the untreated and drug removed lines. No changes were observed in the BFB amplification in Lapatinib drug resistant (LR) and Lapatinib removed (LRDR) cell lines (**Supplementary Figure 13**) The results suggest that unlike ecDNA, BFB structures are more stable and sensitive to targeted therapy. Drug resistance is likely mediated by a change in the population of cells carrying the BFB amplification rather than a change in its structure.

### BFB amplified tumors are more stable compared to ecDNA amplified tumors

We observed significantly longer overall survival in patients with BFB (+)/ecDNA (-) relative to the ecDNA (+) cohort, out to 1100 days (**Figure 5A, P-value 0.02, Log rank test**). Although this difference was no longer seen at longer time points (**Supplementary Figure 14**), these data raise the possibility that ecDNA may be linked to treatment resistance, which usually occurs earlier in the course of the disease. Next, we asked if the BFB structures acquired additional rearrangements (became more complex) as the cancer progressed. In an earlier study, we developed a measure of amplicon complexity based on entropy of the number of genome segments in the focal amplification and the distribution of copy numbers assigned to genome paths extracted by AmpliconArchitect.^36^ The manuscript also showed that patients with esophageal cancer (EAC) and patients with the premalignant Barrett’s esophagus (non-EAC) both carried ecDNA, but the complexity score was higher in the patients with cancer ^36^. Similarly, we also identified BFB amplifications in both non-EAC and EAC patients. Surprisingly, the amplicon complexity in EAC BFBs was lower compared to the BFBs in premalignant non-EAC BFBs (**Figure 5B; p-value 0.001, Mann-Whitney U test**). Together, these results are consistent with BFB amplifications being more stable and less adaptive than ecDNA amplification.

**Figure 5:**
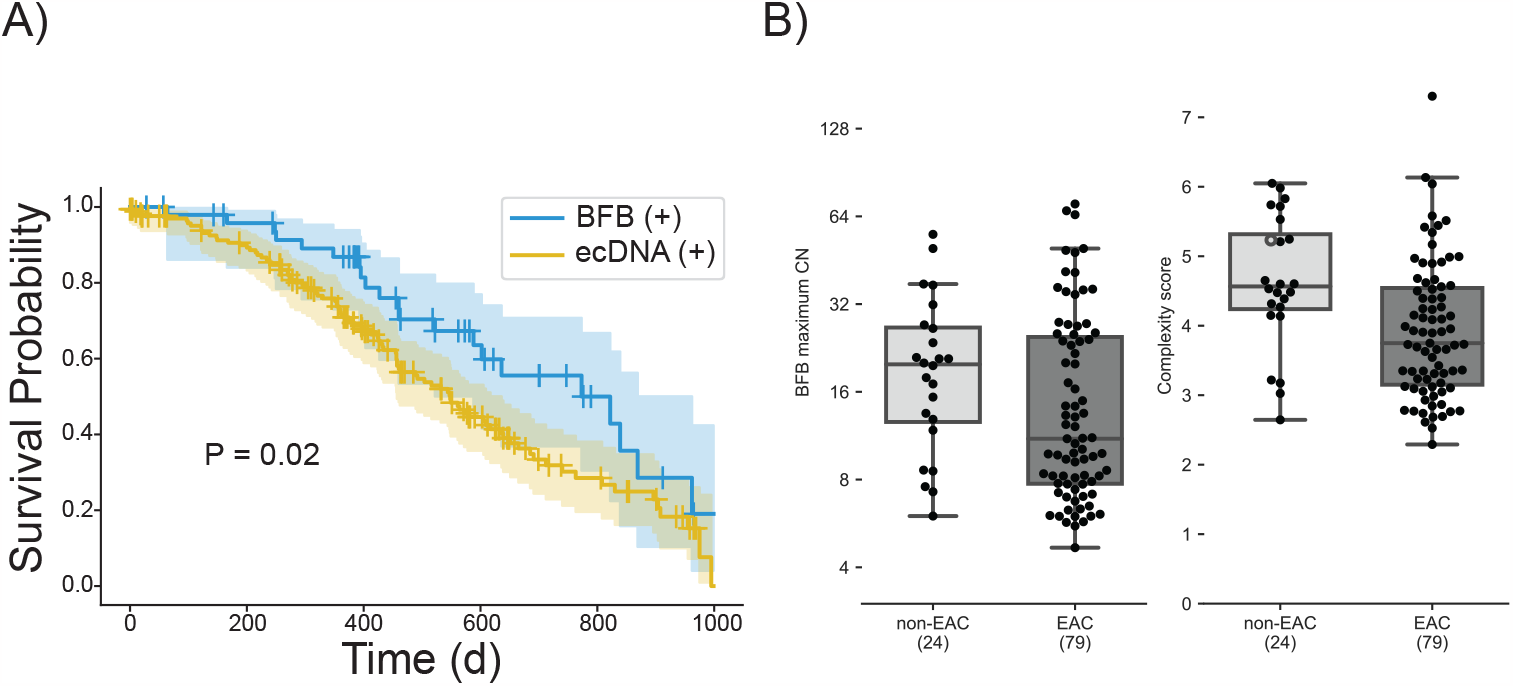
A).Survival outcomes in the first 1000 days (d) for 50 patients with BFB(+) but no ecDNA amplifications (n=50) in their tumors compared to outcomes for ecDNA(+) patients (n=171). P-Value = 0.02 with log rank test. B).Maximum copy number and amplicon complexity scores for BFB amplicons sampled from Barrett’s esophagus (non-EAC) compared to esophageal adenocarcinoma (EAC) patients.

Resistance is possibly acquired through a shift in population of tumor cells carrying specific amplifications. Also, positive selection for specific BFB structures results in reduced heterogeneity over time.

## DISCUSSION

The breakage fusion bridge is an important mechanism of focal oncogene amplification and has previously been observed in many cancers. However, accurate detection of BFB events and characterization of their architecture remains challenging. We found that the typical signature of BFB events–a ladder-like amplification pattern and an excess of foldbacks–may not be sufficient for detection. First, foldback detection is often challenging with short-read sequencing and mapping. This is particularly true when the breaks occur in repetitive or low-complexity regions, or if there is a large distance between the ends of fusing sister chromatids. Second, many non-BFB amplifications, including ecDNA, present foldbacks and sharp ladder-like copy number changes. We resolved the first problem by using longer reads. Specifically, we used optical genome maps which provide megabase length scaffolds that detect foldbacks reliably and also link multiple foldbacks in a single OGM molecule. To resolve the second issue, we developed an algorithm for measuring the likelihood of a BFB amplicon sharing the copy number segmentation and the foldback structures to refine the detection signature. Our method starts by extracting out the processed data into a copy number vector and foldback read vectors. Therefore, it can be extended to nanopore and other long read modalities, simply by changing the initial processing steps, and we plan to do that in future research. As our short read methods use similar abstractions, they also show high precision, but have reduced sensitivity due to missed foldbacks.

The methods we developed here allowed us to systematically explore BFB amplifications in thousands of cancer samples and contrast them with other focal amplifications, including ecDNA. Somewhat surprisingly, we found that the genes amplified by BFB cycles are often also amplified during ecDNA formation and that cancer subtypes with BFB amplification also show ecDNA amplification. However, some genes (e.g. *Myc, CDK4*) are predominantly amplified as ecDNA. Similarly, BFB amplifications are found in most cancers where ecDNA have been observed. However, they are rare in cancers of the brain and central nervous system, where ecDNAs are very common. The reasons for this behavior are not entirely clear. It is plausible that fragile regions initiate BFB formation in specific regions, and not in others. However, our analysis of recurrent BFB amplicons did not support that hypothesis.

While ecDNA and BFB events amplify a similar subset of genes, they are structurally and functionally different. BFB amplicons have nearly twice the span of ecDNA amplicons. However, the localized BFB structure implies that it rarely co-amplifies distant oncogenes, unlike ecDNAs. The increased structural changes and the circularization in ecDNA change their conformation, enabling enhancer hijacking, and the alteration of their TAD boundaries. In contrast, BFBs, which do not have head to tail circularization, or extensive rearrangements, do not create as many novel interactions with distal regions. Increased oncogene expression in BFBs is more likely to be mediated by increase in DNA copy number. Indeed the variability of gene expression after controlling for copy number is lower for BFB amplifications than on ecDNA. The gene expression programs are also markedly different, especially for immune response cells and for checkpoint genes controlling the immune response. These preliminary findings suggest that BFB(+) tumors may not be as immunosuppressive as ecDNA(+) tumors, and might be more amenable to checkpoint inhibition.

Anaphase bridge formation and breakage have been shown to be a precursor to genome instability, including chromothripsis and ecDNA formation^23^. While true, our results also suggest that BFB cycles can lead to a stable amplification step, where only the native chromosome is impacted. We find that the pure BFB amplification studied here is often stable and shows much lower cell to cell heterogeneity compared to ecDNA. This could make BFB amplicons more sensitive to targeted therapy and might anecdotally explain the success of anti-Her2 therapy in HER2+ breast cancers, which are often driven by BFB amplifications^19^.

However, BFB(+) cells can also acquire resistance by population shifts towards cells with alternative amplifications or increased amplification to compensate for the drug. Patient HN137 was initially responsive to anti-EGFR therapy but subsequently developed resistance^34^. While the primary cell line was sensitive to Gefitinib, the untreated metastatic cell line was resistant, and showed an increased copy number for *EGFR* in the BFB amplification. We observed conserved BFB structures that amplified EGFR in both HN137Pri and HN137Met. Surprisingly, HN137Met showed two chromosomal foci, as well as a small number of ecDNA. These early data suggest some plasticity in BFB amplifications and raise the intriguing possibility that ecDNAs mediate the translocation of the BFB structure across chromosomes, or that increasing genomic instability resulted in ecDA formation and translocation of the BFB.

HN137Met, but not HN137Pri, showed *YAP1* amplification on BFB. This likely explains the sensitivity of the Met line to YM155, an inhibitor of BIRC5 and the Yap-Hippo pathway, and shows a shift in the clonality of cell populations between HN137Pri and HN137Met. In the larger cancer genome atlas data, we observed that patients with BFB(+) amplicons, but not ecDNA, had better survival outcomes in the initial period but the advantage was subsequently lost. While these results should be revisited and refined with larger data sets, they support the notion of BFB amplicons being more stable, and less adaptive relative to ecDNA, resulting in delayed onset of resistance.

Nearly 80 years after it was first discovered by Barbara McClintock in irradiated maize cells, the BFB cycle stands firm as an important and distinct mode of focal oncogene amplification in cancer. Identification and reclassification of cancers based on the mechanism of focal amplification is likely to provide more insights into cancer pathology and treatment options.

## METHODS

### Bionano optical genome mapping

Ultra-high molecular weight (UHMW) DNA was isolated from cells using a Bionano Prep SP Blood and Cell Culture DNA Isolation kit (#80042). In brief, about 1 million cells for each sample were lysed and digested in a mixed buffer containing Proteinase K, RNase A, and LBB lysis buffer following the manufacturer’s instructions (Bionano Genomics). A Nanobind Disk was then added to the lysate to bind genomic DNA (gDNA) upon the addition of isopropanol. After washing, the gDNA was eluted and subjected to limited shearing to increase homogeneity by slowly pipetting up and down using standard 200 ul tips.

The gDNA was then equilibrated overnight at room temperature to enhance homogeneity. 2 ul of gDNA aliquot was diluted in Qubit BR buffer and sonicated for 15 min before measuring concentrations with the Qubit dsDNA BR assay kit (Invitrogen Q3285). The UHMW gDNA was ready for labeling when the coefficient of variation of the Qubit reads were less than 0.3. 750ng purified UHMW DNA was fluorescently labeled at the recognition site CTTAAG with the enzyme DLE-1 and subsequently counter-stained using a Bionano Prep DLS Labeling Kit (#80005) following manufacturer’s instructions (Bionano Prep Direct Label and Stain (DLS) Protocol #30206). OGM was performed using a Saphyr platform. Calling of low allele frequency structural variants was performed using the rare variant analysis pipeline (Bionano Solve version 3.6) on molecules ≥ 150kbp in length. The rare variant pipeline enables the detection of SVs occurring at low allelic fractions. Molecules were aligned to the GRCh38 reference, and clusters of molecules (≥3) indicating SVs were used for local assembly. Local consensus assemblies had high accuracy and were used to make final SV calls by realignment to the reference genome. Separately, based on coverage, the pipeline also generated a copy number profile that identified gains and losses. Briefly, molecules were aligned to the GRCh38 reference to create a coverage profile that was then normalized based on OGM controls and scaled against a baseline defined at CN=2 in autosomes (X and Y have a sex chromosome-specific baseline). Putative copy changes were segmented, and calls were generated. Entire chromosomal aneusomies were likewise defined in the CN algorithm.

### Formalization of BFB cycles

Denote a chromosomal arm using consecutive genomic segments A, B, C, D, starting from the centromere and going towards the telomere. The BFB cycle starts with chromosome breakage or a telomere loss, removing segment D. In a pure BFB cycle, where only a single chromosome is implicated, we could see a bridge formation, leading to the di-centric arm 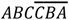, with the bar representing an inversion of the genomic segment. Subsequent breakage between B and A leads to a genome 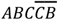, which carries an inverted duplication, and a broken end, allowing for the process to repeat. A small number of BFB cycles lead to a highly rearranged genome. For example,

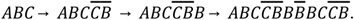

Sampling sequences from the BFB rearranged genome and mapping back to the human reference, we obtain a characteristic *copy-number vector* [1, 6, 4, 0] denoting the copy numbers of segments A through D. Sampling reads from the junction of 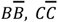, and mapping back to the reference results in *right-fold-back* structures. Similarly, we obtain *left-fold-back* structural variants, corresponding to reads sampled from 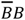.

An abundance of fold-back reads, together with a ladder-like copy number amplification is considered as a signature of BFB. We emphasize, however, that not every structure with an abundance of fold-backs and a ladder like amplification (e.g. 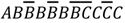) can be explained using BFB cycles, and a more careful exploration is needed.

### Candidate region selection, and parameterization

Conceptually, define an optical genome map (OGM) as a sorted list of numeric values, representing the relative positions of labels on a fragment of DNA. These numeric lists can be generated for any collection of individual OGM molecules, assembled OGM molecules, or from *in silico* predicted label positions on the reference genome. We utilized OGM data from Bionano Genomics, inc. (bionano.com). All OGM samples were pre-processed using the Bionano Solve pipeline. The pipeline aligns and assembles OGM molecules into larger OGM contigs, while also correcting the inter-label distances. We also used Bionano Solve to map the assembled reads to the hg38 genomic reference, call copy numbers and structural variants (SVs). The data from the Bionano Solve pipeline is abstracted through OGM label locations. Each label *i* is covered by *V*_*i*_ molecules. The inferred copy number for label *i* is inferred and represented by *N*_*i*_. Similarly, we denote set of molecules that supports left and right fold-back SVs at label *i* as *F*_*l,i*_ and *F*_*r,i*_, respectively. Thus the data is presented as a tuple (*N, V, F*_*l*_, *F*_*r*_) for each label i. The majority of OGM samples exhibited a coverage range of 100-300x for the diploid chromosome.

### OM2BFB

OM2BFB takes the Bionano Solve output and reconstructs the most likely BFB structure along with the log-likelihood score for reconstruction. better identify foldback reads and copy number profiles. It starts by locating focal amplifications carrying the BFB signature as candidate regions. For each candidate region, it then enumerates multiple possible BFB architectures, modifying methods that were previously developed, including by us, for short read whole genome sequences. Finally, OM2BFB scores each candidate architecture using a novel algorithm to estimate the (negative log-) likelihood of BFB cycle formation. High likelihood (low scoring) reconstructions are output, along with the score as described below. The software is publicly available at https://github.com/siavashre/OM2BFB.

### OM2BFB scoring

Starting with tuples output by the Bionano Solve pipeline, applies some initial filtering steps to identify candidate BFB amplicons. First, candidate labels with a copy number (CN) greater than 3, indicative of amplification, and containing over 10% more foldbacks compared to the average are selected. Subsequently, it determines the span of the amplicon by linking pairs of consecutively selected labels based on two parameters: the distance (D) between the labels and the average copy number (E) of the two labels. Two consecutive regions are clustered if either D < 1.5 Mbp or E>7. These thresholds were empirically determined through the analysis of real BFB datasets. Finally, single linkage clustering was used to identify candidate BFB candidate regions.

To identify BFB structures, recall that a BFB sequence on a genomic region generates a triple (*C, L, R*), described by a CN vector, a ‘left-foldback’ vector, and a ‘right-foldback’ vector, respectively. Note that (*C, L, R*) may not uniquely represent a single BFB sequence, but can also be inferred from the mapped OGM data. OM2BFB computes the ML estimates for (*C, L, R*). Note that the input (*N, V, F*_*l*_, *F*_*r*_) is indexed over labels, while (*C, L, R*) are indexed over genomic segments (each containing a multitude of labels). Therefore, we introduce latent variables (*C*°, *L*° *R*°) referring to observed counts, to estimate

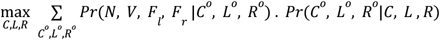

In order to estimate *Pr*(*C*°, *L*°, *R*° |*C, L, R*), we submit the observed vectors to BFB-Finder, which will generate the nearest BFB count vector and multiple candidate BFB sequences, each with their own fold-back vectors. For *n* segments, labeled (*i*, ..., *n*), the Discrepancy between the observed and estimated vectors is computed using

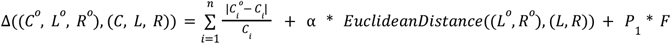

Here, parameter *P*_1_ denotes penalty for foldback discrepancies, and F equals the total number of foldback discrepancies described by

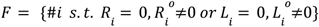

The likelihood of the of (C,L,R) is given by

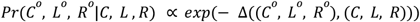

Parameters *P*_1_ = 1, α = 7 were optimized using a grid-search that maximally separated known BFB negative samples from other samples (many but not all of which are BFB positive).

Next, we estimated *Pr*(*N, V, F*_*l*_, *F*_*r*_ |*C*°, *L*°, *R* °), to model the BionanoSolve out which is (*N, V, F*_*l*_, *F*_*r*_) relative to region based copy numbers and fold-backs, given by (*C*°, *L*°, *R*°).

First, we segmented CN vector *N* using the Circular Binary Segmentation (CBS) ^47^. Note that density of labels is not uniform over the reference genome and segment length between two consecutive labels can vary. Therefore, we prepared a weight vector for the CBS algorithm. For each segment, between two consecutive labels, with length larger than 10Kbp we normalized the weight as 1 and for segments with length less than 10 Kbp, the weight was chosen as the segment length in bp divided by 10000. After segmenting with CBS algorithm, we smoothed the result as follows: for each segment *S*_*i*_, if it is a short segment which means that if its length is less than 7 percent of total region length and triplet (*S*_*i*−1_, *S*_*i*_, *S*_*i*+1_) was monotonically increasing or decreasing, segment *S*_*i*_ was merged with the consecutive segment that was closest in copy number. The empirical smoothing removed very short segments associated with sharp increase or decrease in copy numbers. After merging these short segments, the CN vector *C*° was obtained.

Define *Z*_1_ as the expected number of raw molecules that cover a label with copy number 1.

Then, 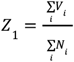, where the indexing is over all labels.

To estimate the CN of each foldback, we divided the number of supported molecules by *Z*_1_. Hence,

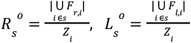

The Discrepancy between the raw input and observed vectors was estimated using:

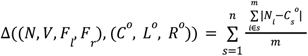

where *m* is number of labels covering by segment *s*. The discrepancy helps estimate the likelihood of (*C* °, *L*°, *R*°) using:

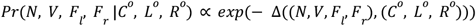

We used a Gibbs sampling approach where in each step, (*C* °_*i*_, *L*°_*i*_, *R*°_*i*_) was sampled for segment *i* using the observed label counts and modifying the observed label counts, merging two adjacent segments, or splitting two segments. At the end of the procedure, the OM2BFB method returns candidate BFB structures that best explain the observed OGM data, along with a score given by:

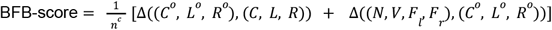

Parameter *c* = 0. 9 was optimized using a grid-search along with previously defined parameters *P*_1_ and α .

OM2BFB is publicly available at https://github.com/siavashre/OM2BFB.

### Visualization

The BFB reconstructions were visualized using a stylized format, as shown in Figure 1B. The axes display genomic coordinates (x-axis) and copy-number (y-axis), with a separate track showing fold-backs. The proposed BFB structure can be read by marking the segments traversed by the blue line, starting from the centromeric end. Red transitions correspond to missing fold-backs required to explain the BFB structure. To avoid huge repetitions of a core structure, OM2BFB depicts only the core structure, along with the multiplicity of repetitions.

### Simulations

BFB molecules were simulated using a custom-developed tool named BFBSimulator, which accepted parameters defining the number of BFB cycles, a chromosome, and start and end positions. Additionally, optional parameters were implemented to allow control over the mean and distribution of BFB segments, along with parameters governing deletion lengths and distribution within the BFB structure and foldback SVs. The tool is publicly available here https://github.com/poloxu/CSE280A_BFBSimulator

Three distinct types of cases were simulated, each representing different complexity levels:

1. **Simple**: Characterized by low segment count, low copy number, no indels, and no deletion in folding regions.
2. **Intermediate**: Regular segment number and copy number were used, along with deletion in folding regions, but no indels were present.
3. **Complex**: This case included regular segment number and copy number, folding region deletions, and indels.

The output from BFBSimulator was formatted as a fasta file. Subsequently, OMSim was employed to simulate optical genome map (OGM) molecules from the fasta output. Parameters recommended by OMSim’s developer were used for simulating the DLE-1 enzyme, but to bypass the assembly of OGM molecules, we simulated long OGM molecules (average length of 10 Mbp) and high accuracy and aligned them to the reference genome. The final step involved using the optical genome map alignment tool FaNDOM^48^ to align the molecules to the reference genome and do the SV calling. In the analysis of OM2BFB, alignments, SV call, and CNV call were essential. Alignments and SV calls from simulated molecules were obtained from FaNDOM and used to create simulated CNV calls. Taking the ground truth CNV values for each genomic segment from the simulated BFB structure we made two different cases. One case segments with exact copy number and contains sharp alterations in segment copy number at segments border, but in the second case, the copy number segmentation boundaries are more gradual, as they appear in real data **(Supplementary Figure 15**).

For simulating BFB negative cases, the focal amplification simulator, ecSimulator^49^(https://github.com/AmpliconSuite/ecSimulator) was employed with different sets of settings to cover simple and complex extrachromosomal DNA (ecDNA) structures in terms of SV rates, with or without duplication inversions. Upon obtaining the fasta file from ecSimulator, the rest of the pipeline, including OGM molecule simulation, contig alignment, and CNV call generation, remained consistent with the process used for BFB positive cases.

### Amplicon classification and BFB detection

We utilized AmpliconClassifier (AC) (version 0.4.11, available from AmpliconSuite at https://github.com/AmpliconSuite/AmpliconClassifier) to identify BFB cycles. AC takes as input the AA breakpoint graph file encoding genomic segment copy numbers and SV breakpoint junctions, as well as the AA cycles file encoding decompositions of the AA graph file into overlapping cyclic and/or non-cyclic paths. Each path and cycle is weighted by the genomic CN it represents. AmpliconClassifier uses multiple heuristics to call BFBs, as described earlier^36^. The salient parts are described below. First AC filters short paths (length <10kbp), paths which significantly overlap low-complexity or repetitive regions, and paths which overlap regions of the genome of low copy number.

AC first assesses non-filtered paths for the presence of BFB cycles using heuristics determined from manual examination of BFB-like focal amplifications in the FHCC cohort and focal amplifications in previous studies^5,24^. AC computes a few relevant statistics: (a) the fraction ‘*f’* of breakpoint graph discordant edges which are foldback, and the paired-ends have a genomic distance < 25kbp. AC next identifies decomposed paths containing foldback junctions between segments, and using all paths computes the set of consecutive segment pairs in the paths where the two boundaries of the segments together form a foldback junction. Each segment pair is assigned its own weight equal to the decomposed copy count of the path. If the proportion of BFB-like segment pairs over all segment pairs in all paths is less than 0.295, then the amplicon is not considered to contain a BFB. Furthermore, if the total weights of pairs which are “distal” (not foldback and > 5kbp jump between endpoints) divided by the total weight of all pairs is greater than 0.5, the amplicon is not considered to contain BFB. Lastly, if the total decomposed CN of all pairs is < 1.5, or if the total number of foldback segment pairs is < 3, or *f* < 0.25, or the decomposed CN weight of all BFB-like paths divided by the CN weight of all paths < 0.6, or the maximum genomic copy number of any region in the candidate BFB region is < 4, the amplicon is not considered to contain a BFB. If the amplicon has not failed any of these criteria, a BFB(+) status is assigned, and the BFB-like cycles (decomposed paths with a BFB foldback) are separated before additional fsCNA detection inside the amplicon region.

### Interphase FISH analysis

In Interphase FISH analysis, the input is an image containing multiple interphase nuclei stained with DAPI and with fluorescent painting of a probed target. The output is a binary segmentation image per probe, with the regions in the image predicted as amplifications are set with value 1 and the background is set with value 0. Our tool returns a binary segmentation image per probe, and each channel (other than DAPI) in the output image is analyzed independently of the other gene probes and other images. The high level steps are as follows:

1. Nuclear segmentation to identify the pixels corresponding to each intact nucleus.
2. Identification and quantification of FISH foci for each nucleus.

These steps are described below.

### FISH Nuclear Segmentation

The chromosomes are unraveled during interphase and occupy much of the nuclear volume. Therefore, the DAPI stain helps separate nuclear regions from the cytoplasmic ones. In patient tissue, however, the nuclei are tightly packed and difficult to resolve into individual nuclei. We applied NuSeT to perform nuclei segmentation^50^. We used a min_score of 0.95, an nms threshold of 0.01, and a scale ratio of 0.3 for all image datasets. For the cell lines with a mix of interphase and metaphase cells (COLO320DM_MYC and HCC827_EGFR), we used a nucleus size threshold of 5000 to prevent individual chromosomes from being classified as nuclei. For the interphase only cell lines (354PDX_ERBB2, 355PDX_ERBB2, 727PDX_ERBB2, Ni8PDX_ERBB2, Ni17PDX_ERBB2, MSTO211H_MYC, COLO320HSR_MYC), we used a nucleus size threshold of 500. Additionally, we applied the min-cut algorithm to convert NuSeT’s binary segmentation output to an instance segmentation. In order to separate neighboring nuclei, for each connected component in the image, we created a pixel graph with 4-connectivity only for pixels with the nuclei binary segmentation value. We looked for pixel centers by convolving the image with a uniform filter, and looked for the minimum number of edges to remove to separate 2 centers in the same connected component in the binary mask. We examined all nuclei with an area larger than 1.25 times the median nuclei size, and separated nuclei with a flow limit of 60.

### Number of FISH spots

To quantify the number of FISH foci, we convolved the original image with a normalized gaussian kernel to determine which pixels have high local intensity. We normalized the gaussian kernel sigma by the median nuclei size for each image. For the median nuclei size of 2500, we used a sampled gaussian kernel with a standard deviation of 3 pixels, and a size of 7 by 7 pixels. After convolving, we applied a threshold of 15 / 255 pixel brightness. Then, to filter out low brightness noise, we have a brightness threshold of 100 / 255 on the original image, and set the minimum spot size to 7 pixels. Finally, to exclude multiple merged nuclei from impacting the FISH spots counts, we removed all nuclei in the top 10 percentile for nuclei area for each sample in the violin plot.

### Test for fragile regions

we investigated the potential influence of positive selection in Breakage-Fusion-Bridge (BFB) regions through a permutation-like test. Seven genomic regions, each containing a minimum of 7 BFBs, were selected. For each region, we divided it into 10 non-overlapping 1Mb windows positioned telomeric to an oncogene. Letting *t*_*i*_ represent the number of BFBs in window *i, s* denoting the total number of BFBs in the region, and *t*_*m*_ representing the maximum number of BFBs observed in a single window. The p-value was calculated as the probability of randomly distributing *s* BFBs into the 10 windows such that some window contained at least *t*_*m*_ BFBs. This probability is equal to

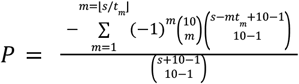

### H3K27ac HiChIP data

We downloaded the hg19-aligned and processed H3K27ac HiChIP for GM12878 from Mumbach et al. 2017^51^ and COLO320DM from Hung et al.2021^39^. We reused the WGS data and cycle structure for COLO320DM using previously reported work^12,39^, where the structure of the *MYC*-amplified ecDNA was fully resolved. We aligned the COLO320DM WGS to hg19 and ran AA (version 1.3.r5) followed by AC (version 0.5.4) to get the list of SVs involving the *MYC*-amplified ecDNA and the BFB-driven focal amplification.

### Distal chromatin interaction identification

We used NeoLoopFinder^44^ version 0.4.3 to search for chromatin interactions on the *MYC*-amplified ecDNA and the BFB-driven focal amplification on chr1 on CN-corrected HiChIP matrices at resolutions 5k and 10k. NeoLoopFinder, by default, computes a genome-wide CN profile and a collection of CN segments from an input contact matrix, and then balances the matrix with a modified ICE procedure by taking the CN segments as input. Given a list of candidate SVs (potentially from other sources, e.g., WGS or OM), it then reconstructs local assemblies representing a chain of one or more SVs from the input list, by shifting or flipping the submatrices according to the coordinates and orientations of the SVs. Therefore, we supplied NeoLoopFinder with a collection of SV breakpoints derived from AA. Additionally, we augmented the assemblies constructed by NeoLooFinder with the collection of local assemblies from the known ecDNA structure in Hung et al. 2021 as follows. Because NeoLoopFinder does not accept assemblies with duplicated segments, we broke the 4.3Mbp ecDNA cycle into all possible longest paths of at least 2 non-overlapping segments. This resulted in 6 distinct assemblies, which were provided as input to NeoLoopFinder to search for chromatin interactions (**Supplementary Table 5**) in addition to the local assemblies constructed above. NeoLoopFinder identified interactions from these HiChIP matrices at different resolutions, and merged the results. It outputs two types of interactions: ‘loops,’ which represent interactions on a single genomic segment, and ‘neo-loops,’ representing interactions on two different genomic segments, brought together by an SV.

As NeoLoopFinder does not support SVs which induce overlapping segments, including fold-back SVs, no neo-loops spanning a fold-back SV were reported. However, we identified loops by providing NeoLoopFinder with a BFB-driven local assembly on chr1 that fully covered the fold-back SVs.

On the control cell line GM12878, whole genome distal chromatin interactions were identified using the method in Salameh et al.^52^ (which was called internally by NeoLoopFinder) and with the same set of resolutions (5k, 10k).

We compared the number of interactions identified by NeoLoopFinder at different distances, normalized by the size of the focal amplifications, for the *MYC*-amplified ecDNA, the BFB-driven amplification and the control cell line GM12878. For GM12878 we included all chromatin interactions and normalized the count at a particular distance by the total genome length. The distribution can be visualized in **Figure 4C**. In order to assess the statistical significance of the differences between the distributions of BFB and ecDNA, we employed the Peacock test^53,54^, a multidimensional version of the Kolmogorov-Smirnov test. This test is specifically designed to analyze random samples defined in two or three dimensions.

### Focal amplification classification from paired-end WGS

We utilized previously published AmpliconArchitect outputs^5^ from TCGA tumor samples and deployed AmpliconClassifier (AC) version 0.4.11 (default settings) to predict presence of BFB in the samples. AC also annotated structures based on gene contents, copy number, and reported an entropy-based complexity score for the focal amplification, using methods described in Luebeck et al.^36^

### Cell culture

Cell lines were purchased from ATCC or Sekisui Xenotech. BT474 was maintained in ATCC Hybri-Care Medium (ATCC, #46-X) with 10% FBS and 1% PSQ. COLO320DM, COLO320HSR and PC3 were cultured in DMEM (Corning, #10-013-CV) with 10% FBS and 1% PSQ. HARA, H460, HCC827, OVCAR3 and SJSA1 were cultured in RPMI-1640 (ATCC modification) (Gibco, #A1049101) with 10% FBS and 1% PSQ. THP1 was cultured in ATCC-formulated RPMI-1640, supplemented with 0.05mM 2-mercaptoethanol and 10% FBS. SNU16M1 was cultured in F12/DMEM supplemented with 10% FBS and 1% PSQ. HCC827 naive, drug resistant (HCC827 ER, HCC827 LR) and drug removed lines (HCC827 ERDR) were cultured in RPMI-1640 (ATCC modification) (Gibco, A1049101) with 10% FBS and 1% PSQ. 3μM erlotinib and 1μM lapatinib were added to HCC827 ER and HCC827 LR respectively to maintain drug resistance. All cell lines were maintained in a 37°C tissue culture incubator supplemented with 5% CO_2_.

### Metaphase FISH

Cell were arrested in mitosis by KaryoMAX Colcemid (Gibco) treatment at 100 ng ml−1 for 4 hours. Cells were washed once in 1X PBS and incubated in 0.075M KCl hypotonic buffer for 20 mins at 37°C. Carnoy fixative (3:1 methanol:glacial acetic acid) were added to fix and wash cells for a total of 3 times. Cells were dropped onto a humidified glass slide and completely air dried. The slides were equilibrated briefly in 2X SSC buffer, followed by ethanol dehydration in ascending ethanol concentrations (70%, 85% then 100%) for 2 mins each. FISH probes diluted in hybridization buffer (Empire Genomics) at 1:6 ratio were added to the sample and sealed with a coverslip. DNA strands were denatured at 75°C for 3 mins, followed by hybridization at 37°C overnight in a dark humidified chamber. Coverslips were removed and samples were washed in 0.4X SSC for 2 mins, followed by 2X SSC 0.1% Tween 20 for 2 mins, and rinsed briefly in 2X SSC. DAPI (50 ng/mL) was used to stain nuclei for 2 mins, and washed briefly in ddH_2_O. Air-dried samples were then mounted with ProLong Diamond Antifade (Invitrogen, #P36931) and cured for at least 4 hours prior to imaging. Images were acquired on a Leica DMi8 widefield microscope on a 63X oil lens.

FISH probes were obtained from Empire Genomics (CCND1, #CCND1-20-GR; CCNE1, #CCNE1-20-RE; FANCG, #FANCG-20-GR; FGFR1, #FGFR1-20-RE; FGFR2, #FGFR2-20-RE; Myc, #MYC-20-GR; Myc-L1, #MYCL1-20-RE; PAK1, #PAK1-20-RE; PDHX, #PDHX-20-RE;

Chromosome 9 Control Probe, #CHR9-10-RE; Chromosome 11 Control Probe, #CHR11-10-AQ), OGT CytoCell (EGFR Amplification Probe, #LPS 003; MDM2 Amplification Probe, #LPS016), Metasystem (XCEP1, #D-0801-050-FI; XCEP10, #D-0810-050-FI) and

Agilent (SureFISH chromosome 19 probe, #G101075-85501).

### Tissue FISH

FFPE slides were deparaffinized by two exchanges of xylene for 5 mins each. The slides were rehydrated in 100% ethanol for 5 mins, followed by 70% ethanol wash for another 5 mins. Slides were briefly rinsed in ddH_2_O and immersed in 0.2N HCl for 20 mins. Antigen retrieval was performed by immersing the slides in 10mM citric acid solution (pH 6.0) and microwaved to reach a temperature at about 90-95°C for 15 mins. Slides were rinsed briefly in 2X SSC and digested with 1μL Proteinase K (NEB, #P8107S) diluted in 100 μL TE buffer at room temperature for 1 min, and washed briefly in ddH_2_O. Slides were then dehydrated in a series of ascending ethanol (70%, 85% and 100%) for 2 mins each. FISH probes diluted in hybridization buffer (Empire Genomics) in 1:6 ratio were applied to the slide. Samples were denatured at 75°C for 3 mins and hybridized at 37°C overnight in a dark humidified chamber. Slides were washed in 0.4X SSC (0.3% IGEPAL) at 40°C for 5 mins twice, followed by another wash in 2X SSC (0.1% IGEPAL) at room temperature for 5 mins. Auto-fluorescence was quenched following the manufacturer’s instructions of the Vector TrueVIEW Autofluorescence Quenching Kit (Vector Laboratories, #SP-8400-15). The slides were stained in DAPI (50 ng/mL) for 10 mins, rinsed twice in 2X SSC and once in ddH_2_O. Air-dried slides were mounted with Prolong Diamond Antifade (Invitrogen, #P36931) and cured overnight at room temperature. Images were acquired on a Leica DMi8 widefield microscope using a 63X oil objective, z-stack images were post-processed using Small Volume Computation Clearing on the LAS X thunder imager prior to generating max projections.

FISH probes were obtained from Empire Genomics (ERBB2, #ERBB2-20-GR; KCTD8, #KCTD8-20-GR; PAX6, #PAX6-20-RE; RP11-1029O18 FISH Probe-Green; Chromosome 4 Control Probe, #CHR4-10-RE; Chromosome 11 Control Probe, #CHR11-10-AQ) and KromaTiD (subCEP CHR 5p Pinpoint FISH Probe, #CEP-0009-C; subCEP CHR 17p Pinpoint FISH Probe, #CEP-0033-D).

### WGS sample and library preparation

gDNA was extracted with the Qiagen DNA Mini Kit following manufacturer’s instructions. 250 ng of gDNA were used to generate sequencing libraries using the NEBNext Ultra II FS DNA Library Prep Kit, following the manufacturer’s protocol to yield the final library with a size distribution between 320-470 bp.

### Bionano optical mapping sample preparation

Cell culture were maintained as described above and a total of 1.5M cells were pelleted in a 1.5mL microfuge tube. The cells were washed twice in 1X PBS and snap-frozen at -80°C prior to shipment to Bionano Genomics for 100x Human Genome Sample Analysis.

## ACKNOWLEDGMENTS

VB, SM, JL, KZ, and BC were supported in part by grants U24CA264379, R01GM114362, and OT2CA278635 from the NIH and CGCATF-2021/100025 from CRUK. GX was supported by a postdoctoral fellowship from the Paul F. Glenn Center for Biology of Aging Research at the Salk Institute. The authors would like to acknowledge Dr. Rebecca Fitzgerald (Cambridge University, UK) and the Seattle Barrett’s study for providing AmpliconArchitect results from Barrett’s esophagus and esophageal cancer samples.

## Fig

**Supplementary Figure 1: Illustration of the Breakage-Fusion-Bridge (BFB) Mechanism leading to Focal Copy Number Amplification**. The BFB mechanism starts with a telomeric break (loss of segment D) that is stabilized by the formation of an anaphase bridge between sister chromatids. Unequal breakage during cytokinesis leads to an inverted duplication genotype ABC→ ABCCB, with a broken end, leading to multiple cycles of bridge formation and breakage until the telomere is recapped, resulting in stable focal amplification with an excess of foldbacks and ladder-like amplification structure.

**Supplementary Figure 2: Distribution of Simulated Cases over the Average Segments Copy Number and Number of Segments**. A scatter plot of the BFB(+) and BFB(-) simulations, characterized by their average segment copy number and the number of segments showing little bias between positive and negative examples. Consistent with prior knowledge, ecDNAs, part of the BFB(-) examples, have higher CN values.

**Supplementary Figure 3: Evaluating the Scoring Metrics for Distinguishing BFB Positive and Negative Cases**. The plot shows the score distribution of 595 BFB(+) and the 191 BFB(-) cases remaining after discarding BFB(-) cases that did not meet the filtering threshold. OM2BFB scoring has 3 components: a copy number score, a segmentation score and a (Euclidean) distance from a candidate BFB. (A) The distribution achieved by the final score, and its performance in terms of the F1 value. (B-D) The distribution and performance after removing one of the 3 components of the score function. (E-H) The distribution and performance after retaining only one of the 3 components of the score function. The results suggest that each component distinctively contributes to improving the performance.

**Supplementary Figure 4: Exemplars of false OM2BFB Predictions in simulated cases**. (Top row) ‘False-negative’ cases that are BFB(+) but were erroneously predicted as BFB(-). The absence of foldback reads (represented in red) reduces the overall score (Left and Middle). Additionally, OM2BFB filters out instances where only one segment remains after segmentation, as those might correspond to ecDNA that originated as BFB (Right). (Bottom row) ‘False-positive’ cases. Non BFB based inverted duplications followed by ecDNA formation sometimes leads to a BFB-like signature. When the number of segments with fold-backs is large (e.g. ≥ 4), the absence of 1-2 foldbacks does not significantly impact the score, resulting in some false positive predictions.

**Supplementary Figure 5: Validation of OM2BFB results with DNA Metaphase FISH**. (A-D) Metaphase FISH images for cancer cell line samples with OM2BFB scores lower than 1.8 all show HSR formation on the native chromosome. (E) A BFB(-) amplification in a non-native chromosome with an OM2BFB score less than 1.8 (false positive call). (F-K) Metaphase FISH images for cancer cell line samples with OM2BFB scores greater than 1.8 showing either HSR formation on non-native chromosomes (panels F-I), or ecDNA (panels J-K). Metaphase FISH images for Colo320DM (chr8 MYC), Colo320HSR (chr8 MYC) and DU145 (chr14 NFKBIA) can be found in the earlier publication^5^.

**Supplementary Figure 6: Validation of OM2BFB results with DNA Interphase FISH**. (A-E) Interphase FISH images for Breast Cancer with Brain Metastases samples with OM2BFB scores lower than 1.8 showing small numbers of distinct foci with low cell to cell heterogeneity.

(F-H) Interphase FISH images for Breast Cancer with Brain Metastases samples with OM2BFB scores greater than 1.8 showing larger numbers of distinct foci with high cell to cell heterogeneity.

**Supplementary Figure 7: BFB amplifications in primary and metastatic cell lines derived from a patient with head and neck cancer**.

**Supplementary Figure 8: Selected metaphase FISH images targeting EGFR in HN137Met**. Left-side panels show EGFR carrying ecDNA in addition to the two chromosomal foci.

**Supplementary Figure 9: Exemplars of discrepancies between OM2BFB predictions on optical genome map data and Amplicon Classifier on whole genome sequencing data**. In the discrepant examples, OM2BFB prediction was BFB(+) (Right, score <1.8), while AC prediction was BFB(-) (Left). (A, B) Missed foldbacks in wgs data (red dashed); (C) Missed foldbacks and the presence of additional translocations lead to a BFB(-) prediction for AC.

**Supplementary Figure 10: Distribution of first-break locations of recurring BFB amplicons**. Panels A-F denote 7 cases with multiple BFB amplifications. The distribution of the most telomeric break to an amplified oncogene is plotted in 10 windows, each of 1Mb (grey tick marks). No 1Mb window is preferentially chosen except for the ERBB2 amplified BFB sites, where a preferential break occurs in the KRTAP region. A preferred (but not statistically significant) window is also seen in BFB sites involving the CCND1 oncogene.

**Supplementary Figure 11: Gene expression of oncogenes vs copy number:**CN of oncogene versus its fold change in RSEM for oncogenes with copy number >4 in the TCGA data set. Each point represents the average fold change of a specific oncogene across all samples containing this amplified oncogene as BFB or ecDNA.

**Supplementary Figure 12: Gene expression comparison for tumor checkpoint genes**. Comparison of raw gene expression count for tumor checkpoint genes between ecDNA(+) and BFB(+) samples in TCGA dataset. Statistical significance was determined using the rank-sum test.

**Supplementary Figure 13: HCC827-LRDR**. Targeted Therapy Resistance of the HCC827 Cell Line continuing EGFR amplified within a BFB event. The copy number and the proportion of cells carrying the BFB signal are restored after drug removal (LRDR) in the Lapatinib drug resistant cell line along with metaphase FISH images.

**Supplementary Figure 14: Survival rate**. The survival outcomes for 50 patients with BFB but no ecDNA amplifications in their tumors compared to outcomes for 171 patients with ecDNA amplifications. BFB(+) individuals have better outcomes initially (first 1100 days).

**Supplementary Figure 15: CNV call simulation**. Illustrations of two different cases of CNV call simulation. The left panel represents segments of CN without any added noise, resulting in clear and distinct boundaries. In contrast, the right panel includes added noise, making the copy number segmentation boundaries more gradual and less defined.

## Supplementary Table Captions

**Supplementary Table 1:** Cell line analysis using OM2BFB and FISH in both Metaphase and Interphase experiments.

**Supplementary Table 2:** Sample analysis of TCGA, CCLE, and BE using Amplicon Classifier.

**Supplementary Table 3:** Recurrently amplified genomic regions via BFB mechanism at least 7 times with corresponding P-values from BFB distributions calculated with permutation-like test.

**Supplementary Table 4:** Oncogene amplification frequencies via BFB or ecDNA.

**Supplementary Table 5:** NeoLoopFinder assemblies obtained from Amplicon Architect.

## REFERENCES

1. Davoli, T., Uno, H., Wooten, E. C. & Elledge, S. J. Tumor aneuploidy correlates with markers of immune evasion and with reduced response to immunotherapy. Science 355, eaaf8399 (2017).

2. Krijgsman, O., Carvalho, B., Meijer, G. A., Steenbergen, R. D. M. & Ylstra, B. Focal chromosomal copy number aberrations in cancer—Needles in a genome haystack. Biochim. Biophys. Acta BBA - Mol. Cell Res. 1843, 2698–2704 (2014).

3. Zhao, X.-K. et al. Focal amplifications are associated with chromothripsis events and diverse prognoses in gastric cardia adenocarcinoma. Nat. Commun. 12, 6489 (2021).

4. Turner, K. M. et al. Extrachromosomal oncogene amplification drives tumour evolution and genetic heterogeneity. Nature 543, 122–125 (2017).

5. Kim, H. et al. Extrachromosomal DNA is associated with oncogene amplification and poor outcome across multiple cancers. Nat. Genet. 52, 891–897 (2020).

6. Hamkalo, B. A., Farnham, P. J., Johnston, R. & Schimke, R. T. Ultrastructural features of minute chromosomes in a methotrexate-resistant mouse 3T3 cell line. Proc. Natl. Acad. Sci. 82, 1126–1130 (1985).

7. Von Hoff, D. D., Forseth, B., Clare, C. N., Hansen, K. L. & VanDevanter, D. Double minutes arise from circular extrachromosomal DNA intermediates which integrate into chromosomal sites in human HL-60 leukemia cells. J. Clin. Invest. 85, 1887–1895 (1990).

8. Paulsen, T., Kumar, P., Koseoglu, M. M. & Dutta, A. Discoveries of Extrachromosomal Circles of DNA in Normal and Tumor Cells. Trends Genet. 34, 270–278 (2018).

9. Bafna, V. & Mischel, P. S. Extrachromosomal DNA in Cancer. Annu. Rev. Genomics Hum. Genet. 23, 29–52 (2022).

10. Corces, M. R. et al. The chromatin accessibility landscape of primary human cancers. Science 362, eaav1898 (2018).

11. Morton, A. R. et al. Functional Enhancers Shape Extrachromosomal Oncogene Amplifications. Cell 179, 1330–1341.e13 (2019).

12. Wu, S. et al. Circular ecDNA promotes accessible chromatin and high oncogene expression. Nature 575, 699–703 (2019).

13. Koche, R. P. et al. Extrachromosomal circular DNA drives oncogenic genome remodeling in neuroblastoma. Nat. Genet. 52, 29–34 (2020).

14. Kaufman, R. J., Brown, P. C. & Schimke, R. T. Amplified dihydrofolate reductase genes in unstably methotrexate-resistant cells are associated with double minute chromosomes. Proc. Natl. Acad. Sci. U. S. A. 76, 5669–5673 (1979).

15. Nathanson, D. A. et al. Targeted Therapy Resistance Mediated by Dynamic Regulation of Extrachromosomal Mutant EGFR DNA. Science 343, 72–76 (2014).

16. McClintock, B. The Behavior in Successive Nuclear Divisions of a Chromosome Broken at Meiosis. Proc. Natl. Acad. Sci. 25, 405–416 (1939).

17. McClintock, B. THE STABILITY OF BROKEN ENDS OF CHROMOSOMES IN ZEA MAYS. Genetics 26, 234–282 (1941).

18. Gisselsson, D. et al. Chromosomal breakage-fusion-bridge events cause genetic intratumor heterogeneity. Proc. Natl. Acad. Sci. U. S. A. 97, 5357–5362 (2000).

19. Marotta, M. et al. Palindromic amplification of the ERBB2 oncogene in primary HER2-positive breast tumors. Sci. Rep. 7, 1–12 (2017).

20. Ferrari, A. et al. A whole-genome sequence and transcriptome perspective on HER2-positive breast cancers. Nat. Commun. 7, (2016).

21. Difilippantonio, M. J. et al. Evidence for replicative repair of DNA double-strand breaks leading to oncogenic translocation and gene amplification. J. Exp. Med. (2002) doi:10.1084/jem.20020851.

22. Shimizu, N., Shingaki, K., Kaneko-Sasaguri, Y., Hashizume, T. & Kanda, T. When, where and how the bridge breaks: anaphase bridge breakage plays a crucial role in gene amplification and HSR generation. Exp. Cell Res. 302, 233–243 (2005).

23. Umbreit, N. T. et al. Mechanisms generating cancer genome complexity from a single cell division error. Science 368, (2020).

24. Deshpande, V. et al. Exploring the landscape of focal amplifications in cancer using AmpliconArchitect. Nat. Commun. 10, 392 (2019).

25. AmpliconSuite · GitHub. https://github.com/AmpliconSuite.

26. Raeisi Dehkordi, S. GitHub - siavashre/OM2BFB: OM2BFB a tool for detecting breakage fusion bridge cycles with OGM technology. https://github.com/siavashre/OM2BFB (2023).

27. Zakov, S., Kinsella, M. & Bafna, V. An algorithmic approach for breakage-fusion-bridge detection in tumor genomes. Proc. Natl. Acad. Sci. U. S. A. 110, 5546–5551 (2013).

28. Zakov, S. & Bafna, V. Reconstructing Breakage Fusion Bridge Architectures Using Noisy Copy Numbers. J. Comput. Biol. 22, 577–594 (2015).

29. Greenman, C. D., Cooke, S. L., Marshall, J., Stratton, M. R. & Campbell, P. J. Modeling the evolution space of breakage fusion bridge cycles with a stochastic folding process. J. Math. Biol. 72, 47–86 (2016).

30. Luebeck, J. ecSimulator. (2023).

31. Tanaka, H. et al. Intrastrand annealing leads to the formation of a large DNA palindrome and determines the boundaries of genomic amplification in human cancer. Mol. Cell. Biol. 27, 1993–2002 (2007).

32. Ni, J. et al. Combination inhibition of PI3K and mTORC1 yields durable remissions in orthotopic patient-derived xenografts of HER2-positive breast cancer brain metastases. Nat. Med. 22, 723 (2016).

33. Ni, J. et al. p16INK4A-deficiency predicts response to combined HER2 and CDK4/6 inhibition in HER2+ breast cancer brain metastases. Nat. Commun. 13, 1473 (2022).

34. Chia, S. et al. Phenotype-driven precision oncology as a guide for clinical decisions one patient at a time. Nat. Commun. 8, 435 (2017).

35. Shoshani, O. et al. Chromothripsis drives the evolution of gene amplification in cancer. Nature 591, 137–141 (2021).

36. Luebeck, J. et al. Extrachromosomal DNA in the cancerous transformation of Barrett’s oesophagus. Nature 616, 798–805 (2023).

37. Cancer Cell Line Encyclopedia (CCLE). https://sites.broadinstitute.org/ccle/.

38. Suzuki, R. et al. The fragility of a structurally diverse duplication block triggers recurrent genomic amplification. Nucleic Acids Res. 49, 244–256 (2021).

39. Hung, K. L. et al. EcDNA hubs drive cooperative intermolecular oncogene expression. bioRxiv 2020.11.19.390278 https://www.biorxiv.org/content/10.1101/2020.11.19.390278v1 (2021) xdoi:10.1101/2020.11.19.390278.

40. Zhu, Y. et al. Oncogenic extrachromosomal DNA functions as mobile enhancers to globally amplify chromosomal transcription. Cancer Cell 39, 694–707.e7 (2021).

41. Lee, J. J.-K. et al. ERα-associated translocations underlie oncogene amplifications in breast cancer. Nature 1–9 (2023) doi:10.1038/s41586-023-06057-w.

42. Helmsauer, K. et al. Enhancer hijacking determines extrachromosomal circular MYCN amplicon architecture in neuroblastoma. Nat. Commun. 2020 111 11, 1–12 (2020).

43. Turner, K. M. et al. Extrachromosomal oncogene amplification drives tumour evolution and genetic heterogeneity. Nature 543, (2017).

44. Wang, X. et al. Genome-wide detection of enhancer-hijacking events from chromatin interaction data in rearranged genomes. Nat. Methods 18, 661–668 (2021).

45. Wu, T. et al. Extrachromosomal DNA formation enables tumor immune escape potentially through regulating antigen presentation gene expression. Sci. Rep. 12, 3590 (2022).

46. Lange, J. T. et al. The evolutionary dynamics of extrachromosomal DNA in human cancers. Nat. Genet. 54, 1527–1533 (2022).

47. Olshen, A. B., Venkatraman, E. S., Lucito, R. & Wigler, M. Circular binary segmentation for the analysis of array-based DNA copy number data. Biostatistics 5, 557–572 (2004).

48. Raeisi Dehkordi, S., Luebeck, J. & Bafna, V. FaNDOM: Fast nested distance-based seeding of optical maps. Patterns 2, 100248 (2021).

49. Luebeck, J. et al. AmpliconReconstructor integrates NGS and optical mapping to resolve the complex structures of focal amplifications. Nat. Commun. 11, 4374 (2020).

50. Yang, L. et al. NuSeT: A deep learning tool for reliably separating and analyzing crowded cells. PLOS Comput. Biol. 16, e1008193 (2020).

51. Mumbach, M. R. et al. Enhancer connectome in primary human cells identifies target genes of disease-associated DNA elements. Nat. Genet. 49, 1602–1612 (2017).

52. Salameh, T. J. et al. A supervised learning framework for chromatin loop detection in genome-wide contact maps. Nat. Commun. 11, 3428 (2020).

53. Fasano, G. & Franceschini, A. A multidimensional version of the Kolmogorov–Smirnov test. Mon. Not. R. Astron. Soc. 225, 155–170 (1987).42.ndtest. https://github.com/syrte/ndtest

