## Supplementary Figures for "Breakage fusion bridge cycles drive high oncogene copy number, but not intratumoral genetic heterogeneity or rapid cancer genome change"

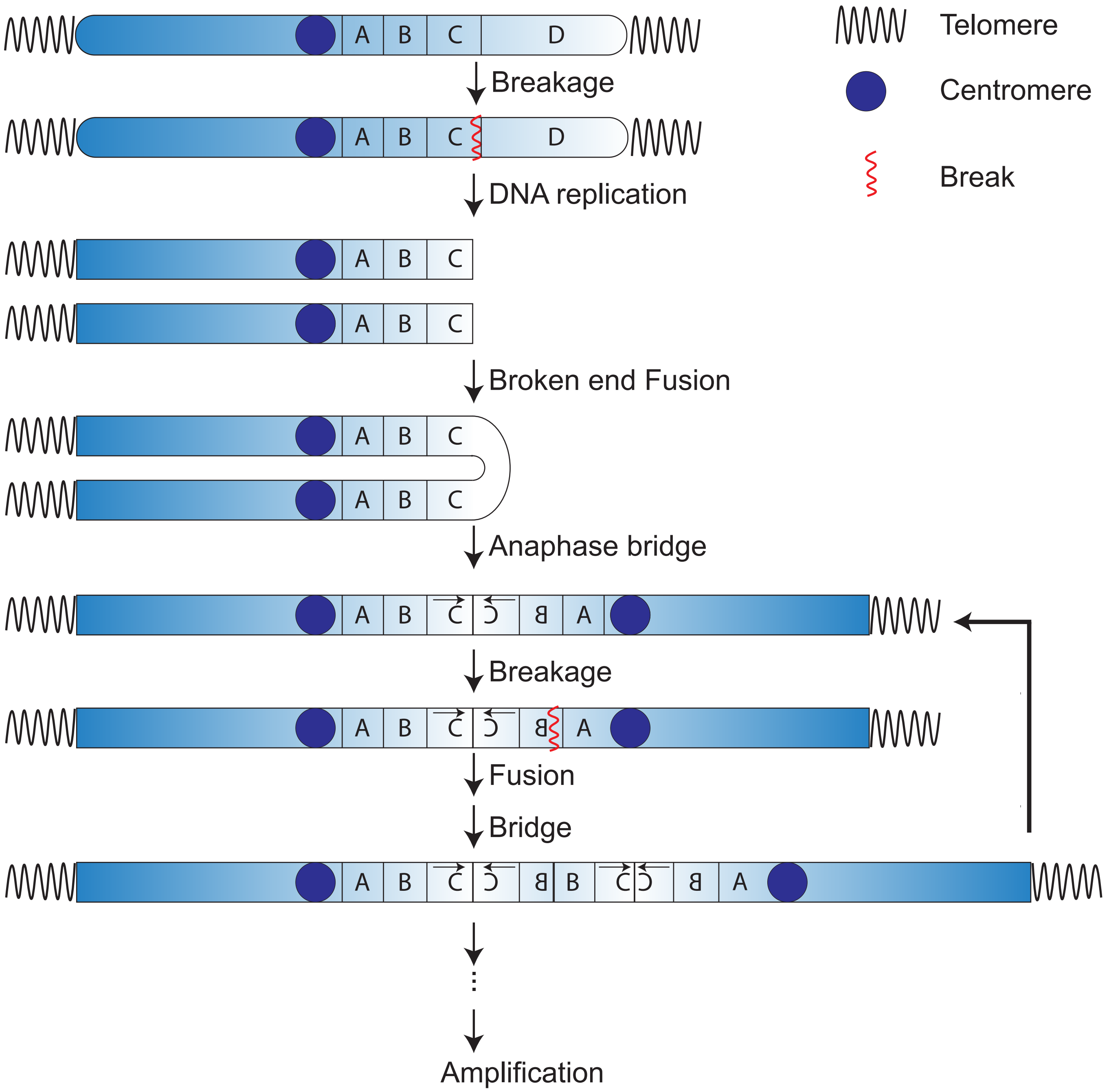

Telomere
Centromere
Break
A B C D
Breakage
A B C D
DNA replication
A B C
A B C
Broken end Fusion
A B C
A B C
Anaphase bridge
A B C A
Breakage
A B C A
Fusion
Bridge
A B C B C A
Amplification

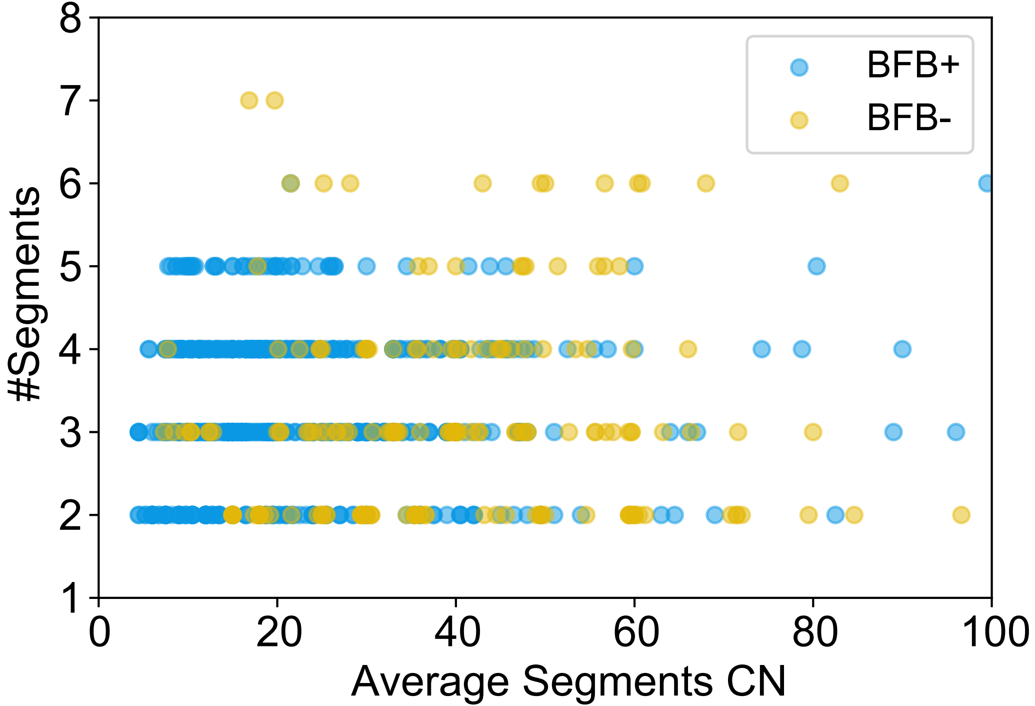

8
7
6
5
4
3
2
1
#Segments
0
20
40
60
80
100
Average Segments CN
BFB+
BFB-

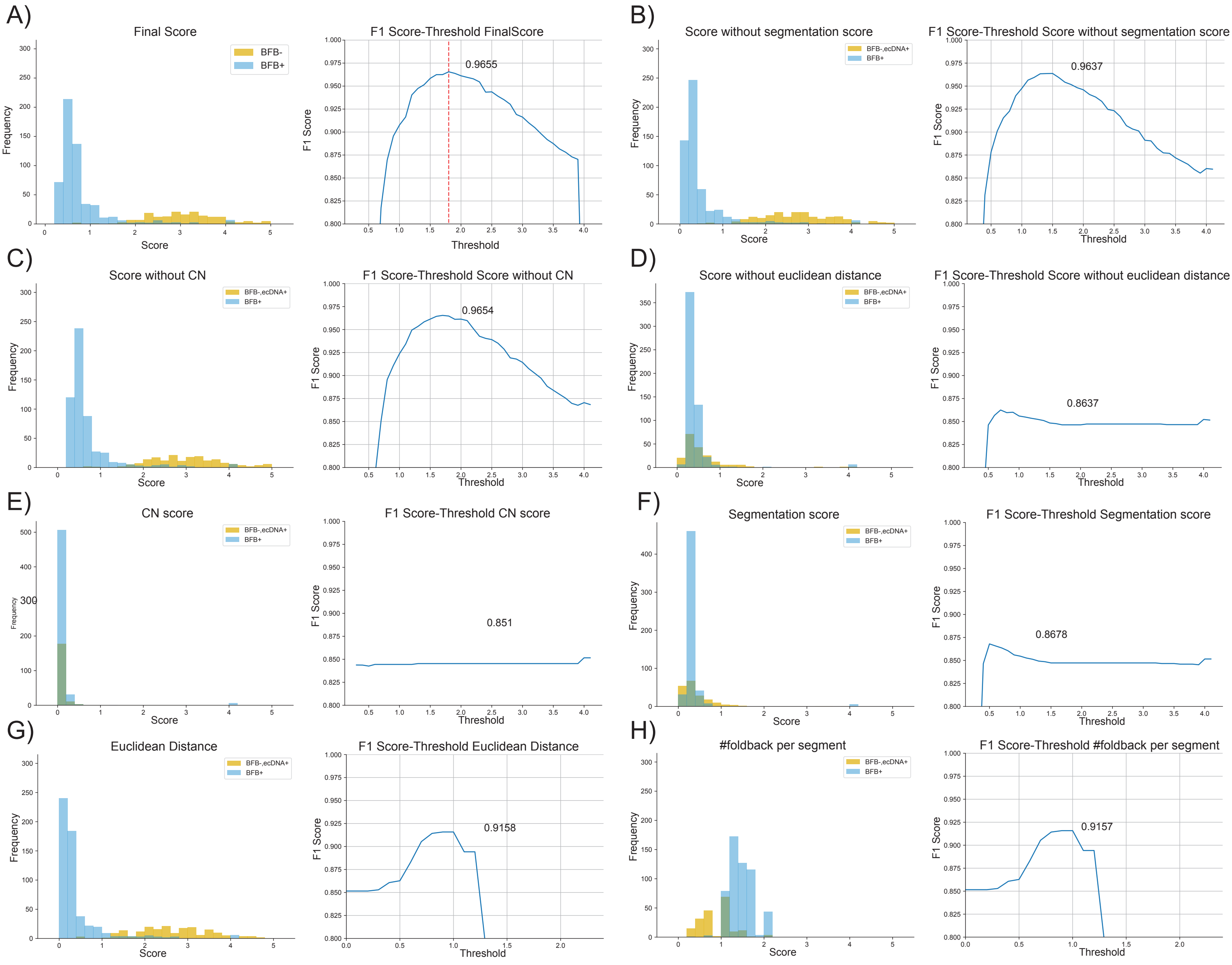
A)
Final Score
BFB-
BFB+
Frequency
Score
F1 Score-Threshold FinalScore
0.9655
F1 Score
Threshold
B)
Score without segmentation score
BFB-,ecDNA+
BFB+
F1 Score-Threshold Score without segmentation score
0.9637
C)
Score without CN
F1 Score-Threshold Score without CN
0.9654
D)
Score without euclidean distance
F1 Score-Threshold Score without euclidean distance
0.8637
E)
CN score
F1 Score-Threshold CN score
0.851
F)
Segmentation score
F1 Score-Threshold Segmentation score
0.8678
G)
Euclidean Distance
F1 Score-Threshold Euclidean Distance
0.9158
H)
#foldback per segment
F1 Score-Threshold #foldback per segment
0.9157

### False Negative

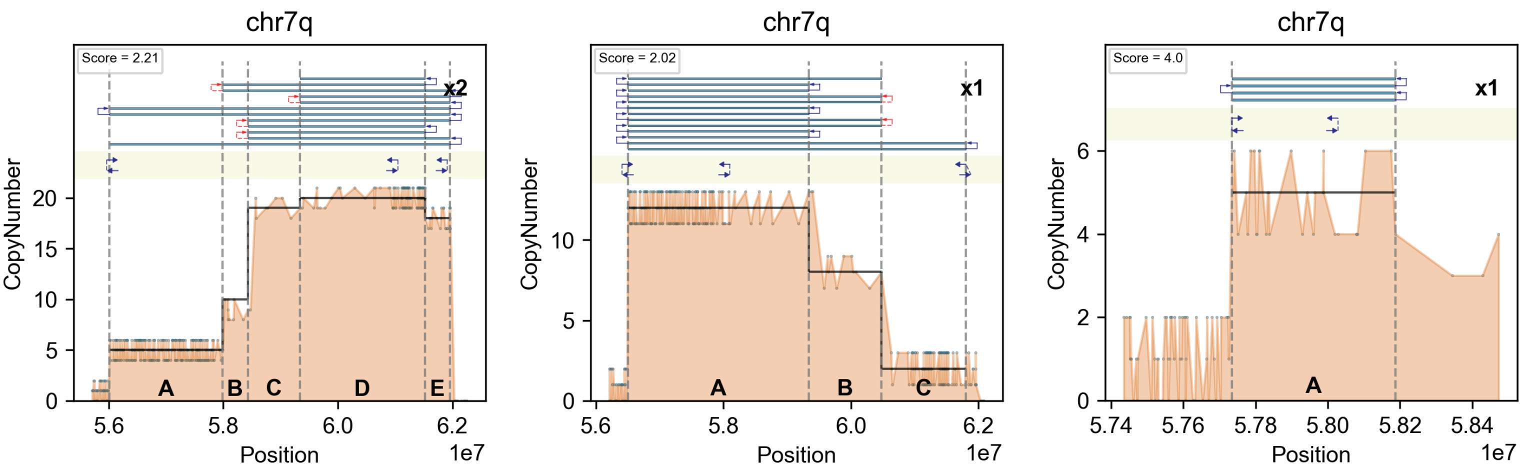


### False Positive

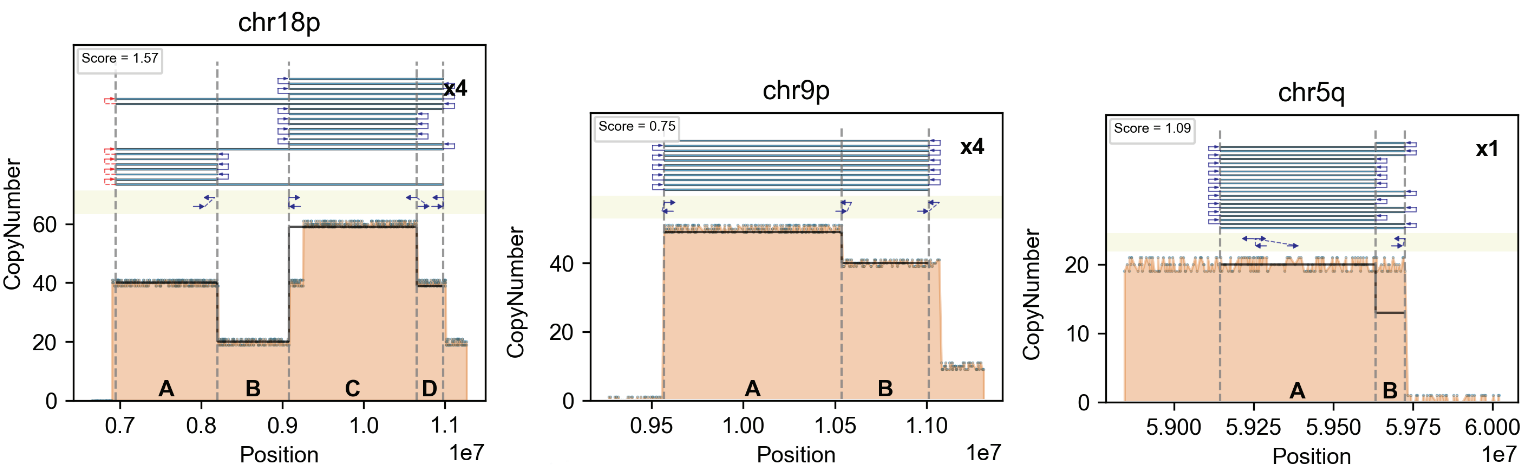

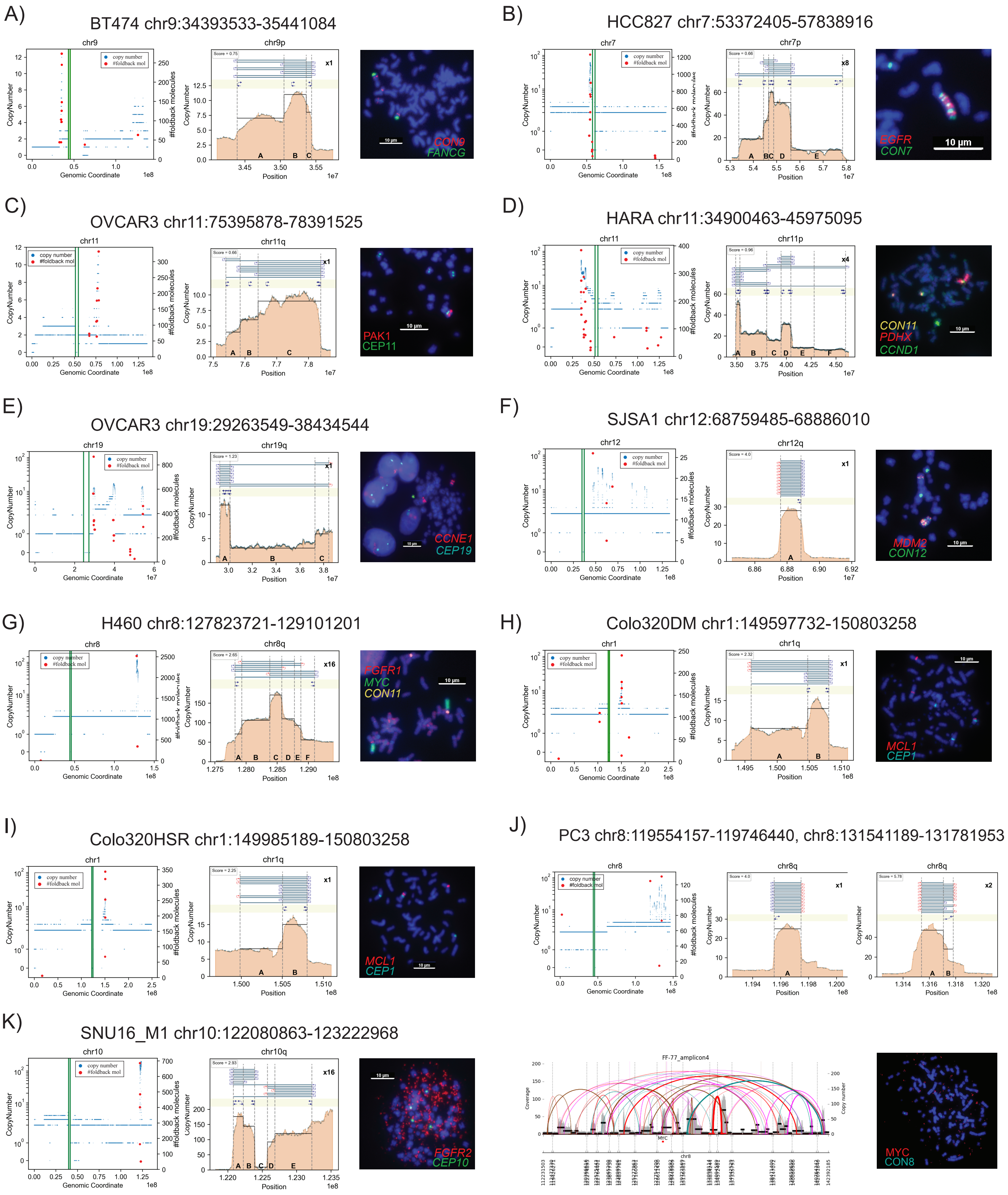
A)
BT474 chr9:34393533-35441084
chr9
chr9p
copy number
#foldback mol
CopyNumber
#foldback molecules
Genomic Coordinate
Position
Score = 0.75
x1
A
B
C
CON9
FANCG
10 μm
B)
HCC827 chr7:53372405-57838916
chr7
chr7p
Score = 0.66
x8
A
BC
D
E
EGFR
CON7
10 μm
C)
OVCAR3 chr11:75395878-78391525
chr11
chr11q
Score = 0.66
x1
A
B
C
PAK1
CEP11
10 μm
D)
HARA chr11:34900463-45975095
chr11
chr11p
Score = 0.96
x4
A
B
C
D
E
F
CON11
PDHX
CCND1
10 μm
E)
OVCAR3 chr19:29263549-38434544
chr19
chr19q
Score = 1.23
x1
A
B
C
CCNE1
CEP19
10 μm
F)
SJSA1 chr12:68759485-68886010
chr12
chr12q
Score = 4.0
x1
A
MDM2
CON12
10 μm
G)
H460 chr8:127823721-129101201
chr8
chr8q
Score = 2.65
x16
A
B
C
D
E
F
FGFR1
MYC
CON11
10 μm
H)
Colo320DM chr1:149597732-150803258
chr1
chr1q
Score = 2.32
x1
A
B
MCL1
CEP1
10 μm
I)
Colo320HSR chr1:149985189-150803258
chr1
chr1q
Score = 2.25
x1
A
B
MCL1
CEP1
10 μm
J)
PC3 chr8:119554157-119746440, chr8:131541189-131781953
chr8
chr8q
Score = 4.0
x1
Score = 5.78
x2
A
B
K)
SNU16_M1 chr10:122080863-123222968
chr10
chr10q
Score = 2.93
x16
A
B
C
D
E
FGFR2
CEP10
10 μm
FF-77_amplicon4
Coverage
Copy number
MYC
chr8
MYC
CON8

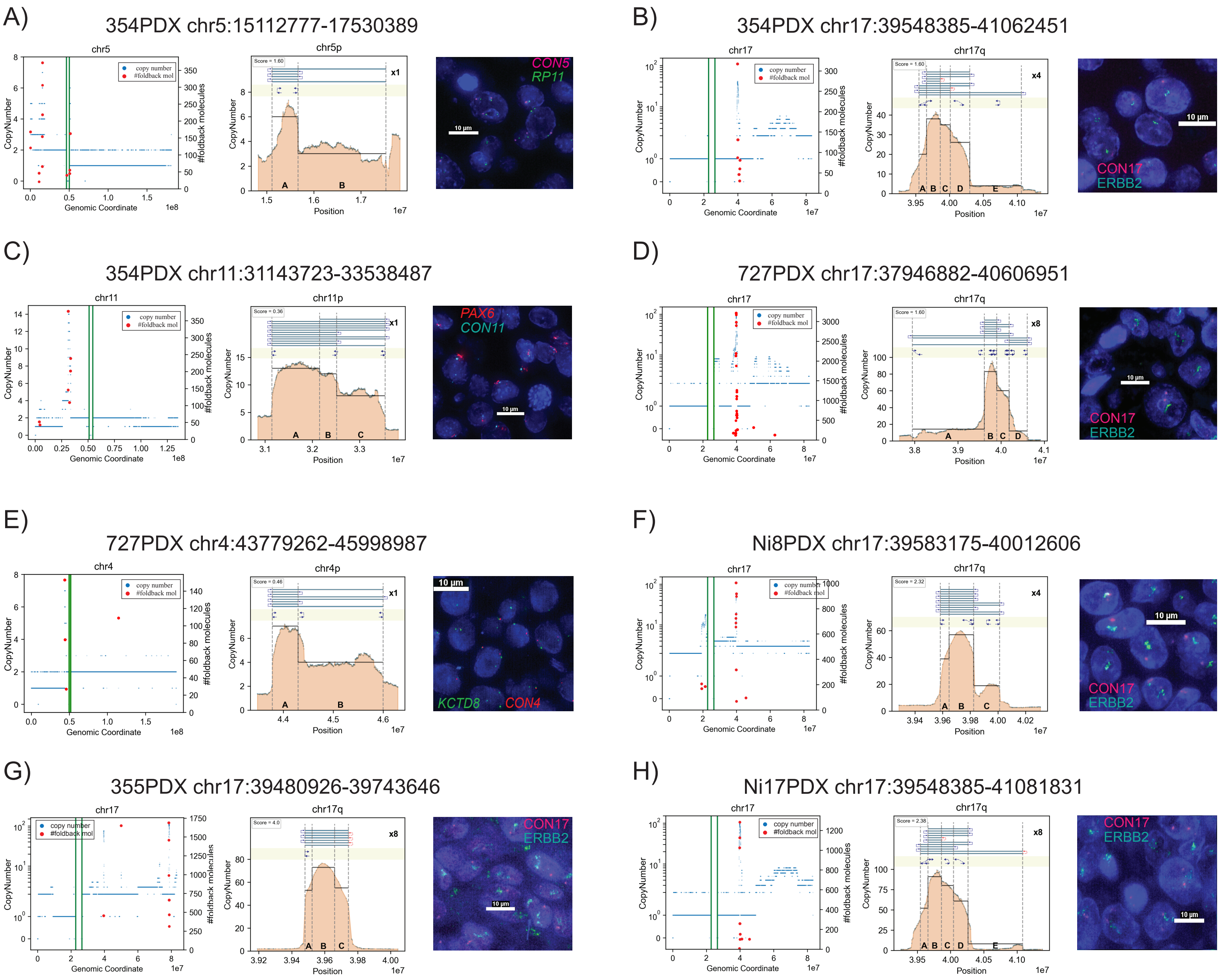
A)
354PDX chr5:15112777-17530389
chr5
chr5p
CON5
RP11
B)
354PDX chr17:39548385-41062451
chr17
chr17q
CON17
ERBB2
C)
354PDX chr11:31143723-33538487
chr11
chr11p
PAX6
CON11
D)
727PDX chr17:37946882-40606951
chr17
chr17q
CON17
ERBB2
E)
727PDX chr4:43779262-45998987
chr4
chr4p
KCTD8
CON4
F)
Ni8PDX chr17:39583175-40012606
chr17
chr17q
CON17
ERBB2
G)
355PDX chr17:39480926-39743646
chr17
chr17q
CON17
ERBB2
H)
Ni17PDX chr17:39548385-41081831
chr17
chr17q
CON17
ERBB2
copy number
#foldback mol
CopyNumber
#foldback molecules
Genomic Coordinate
Position
10 µm

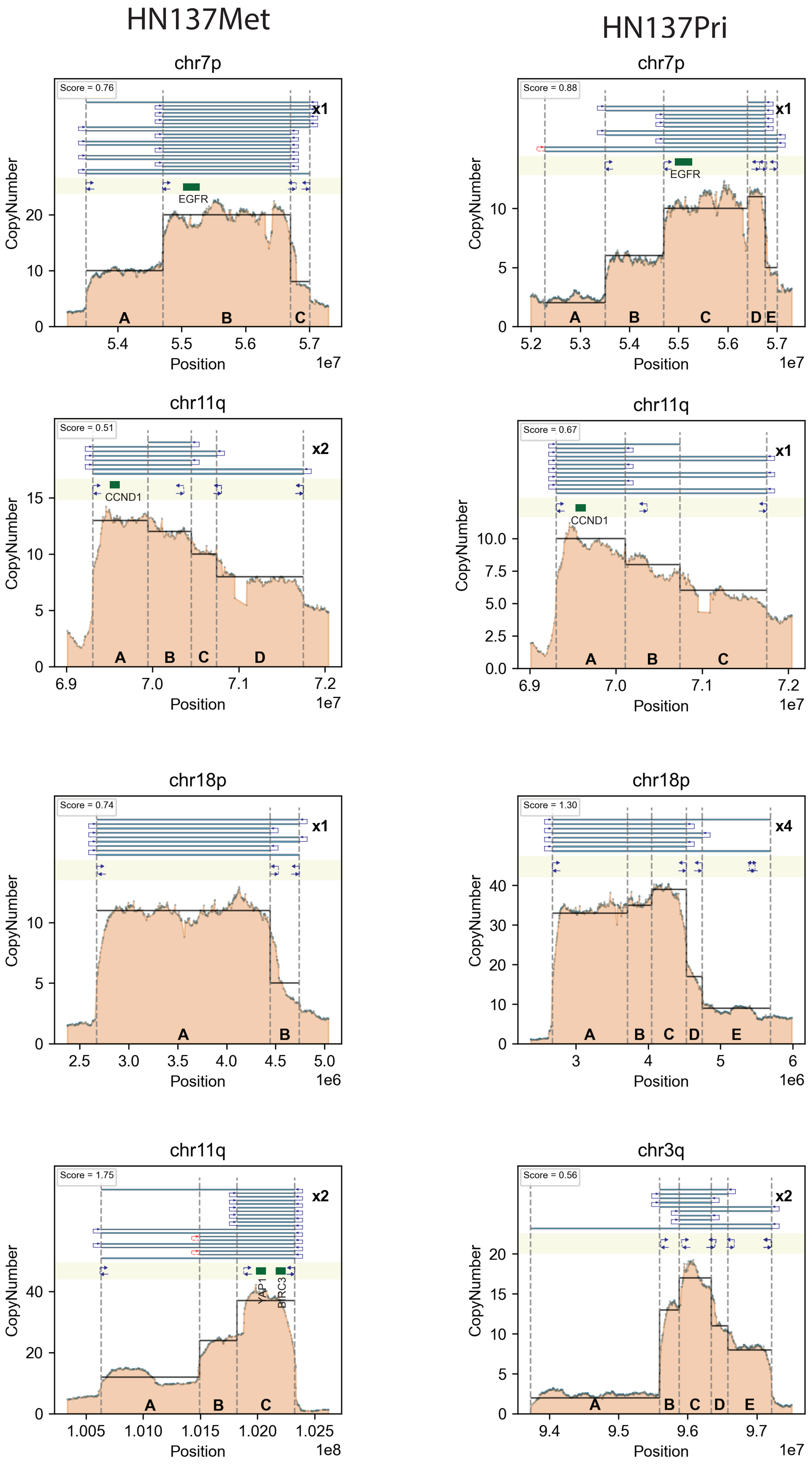
HN137Met
HN137Pri
chr7p
Score = 0.76
x1
EGFR
CopyNumber
Position
1e7
A
B
C
chr7p
Score = 0.88
x1
EGFR
A
B
C
D
E
chr11q
Score = 0.51
x2
CCND1
A
B
C
D
chr11q
Score = 0.67
x1
CCND1
A
B
C
chr18p
Score = 0.74
x1
A
B
1e6
chr18p
Score = 1.30
x4
A
B
C
D
E
chr11q
Score = 1.75
x2
YAP1
BIRC3
A
B
C
1e8
chr3q
Score = 0.56
x2
A
B
C
D
E

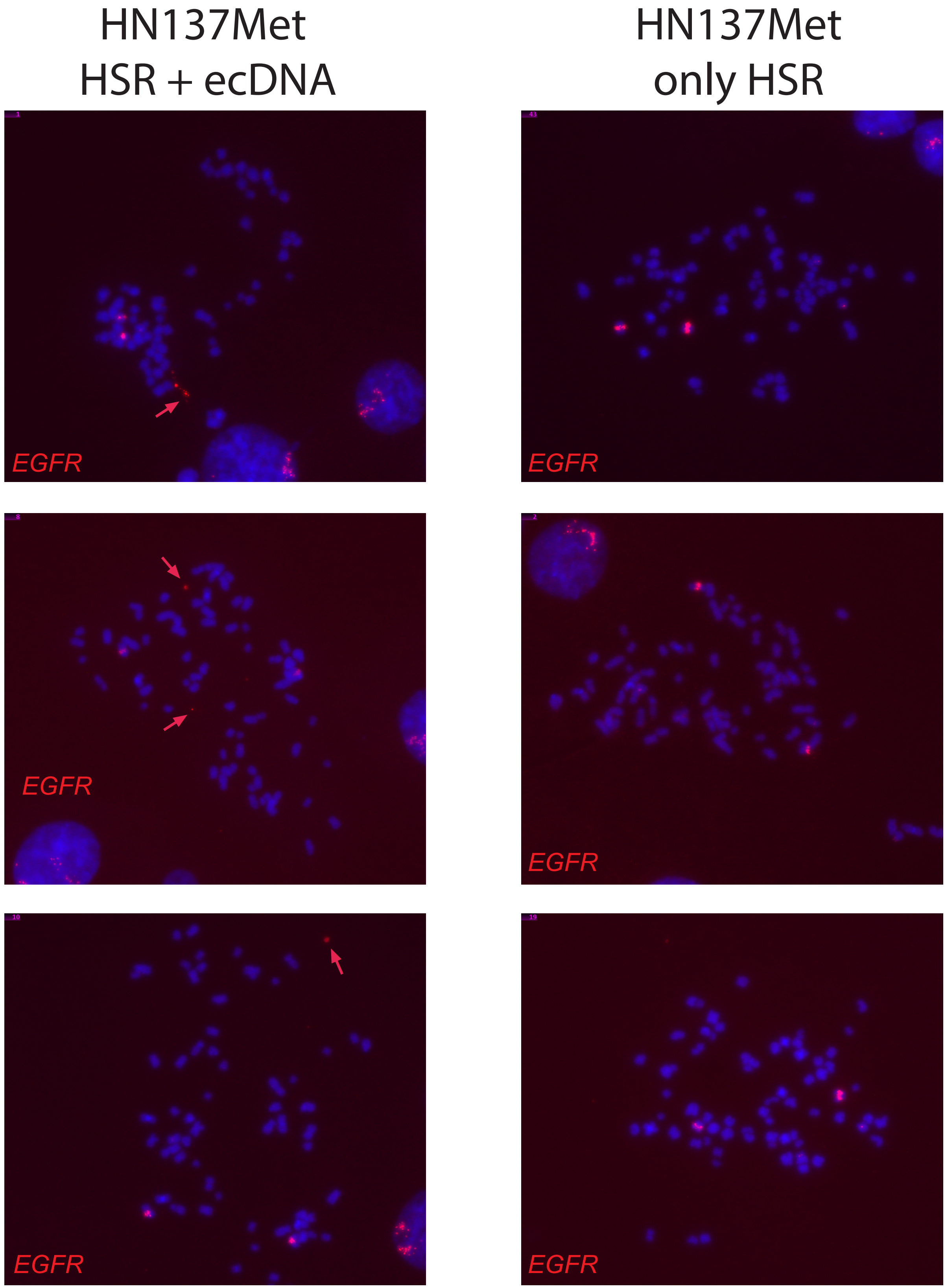
HN137Met
HSR + ecDNA
HN137Met
only HSR
EGFR
EGFR
EGFR
EGFR
EGFR
EGFR

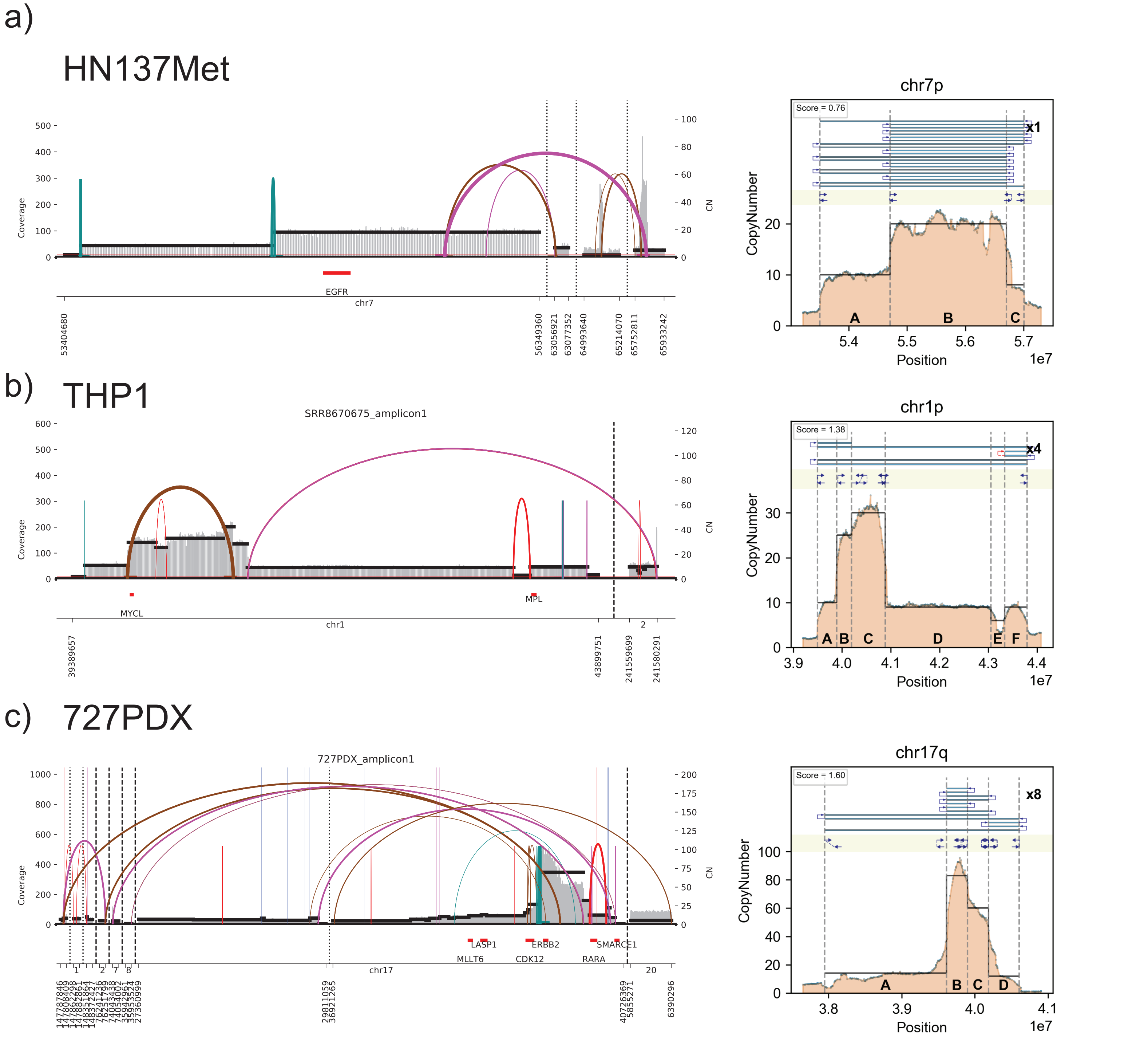
a)
HN137Met
Coverage
CN
EGFR
chr7
53404680
56349360
63056921
63077352
64993640
65214070
65752811
65933242
chr7p
Score = 0.76
x1
CopyNumber
A
B
C
Position
1e7
b)
THP1
SRR8670675_amplicon1
Coverage
CN
MYCL
MPL
chr1
39389657
43899751
241559699
241580291
chr1p
Score = 1.38
x4
CopyNumber
A
B
C
D
E
F
Position
1e7
c)
727PDX
727PDX_amplicon1
Coverage
CN
LASP1
MLLT6
ERBB2
CDK12
SMARCE1
RARA
chr17
29811059
36921265
40726369
5855271
6390296
chr17q
Score = 1.60
x8
CopyNumber
A
B
C
D
Position
1e7

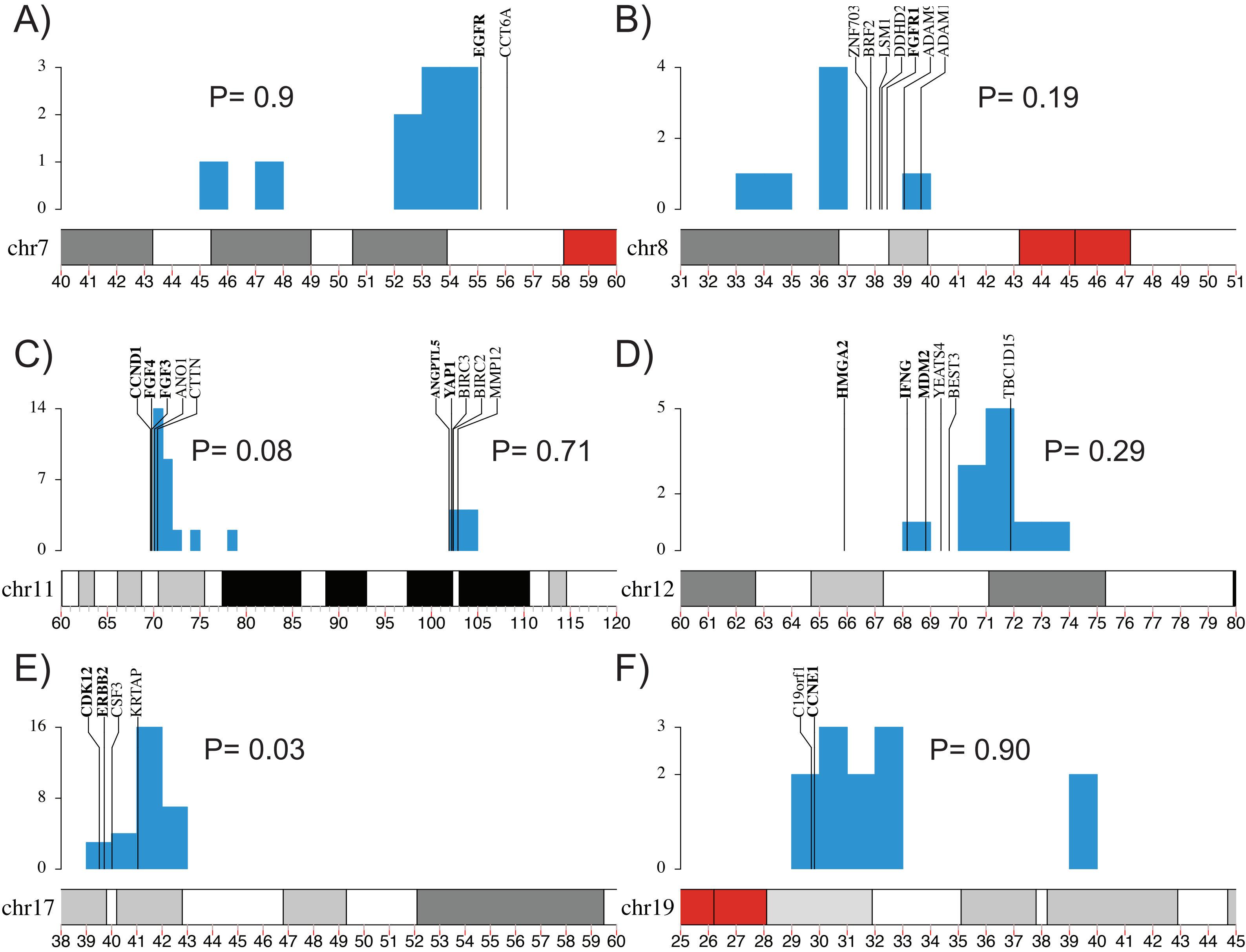

A)
EGFR
CCT6A
P= 0.9
chr7
B)
ZNF703
BRF2
LSM1
DDHD2
FGFR1
P= 0.19
chr8
C)
CCND1
FGF4
FGF3
ANO1
CTTN
ANGPTL5
YAP1
BIRC3
BIRC2
MMP12
P= 0.08
P= 0.71
chr11
D)
HMGA2
IFNG
MDM2
YEATS4
BEST3
TBC1D15
P= 0.29
chr12
E)
CDK12
ERBB2
CSF3
KRTAP
P= 0.03
chr17
F)
C19orf1
CCNE1
P= 0.90
chr19

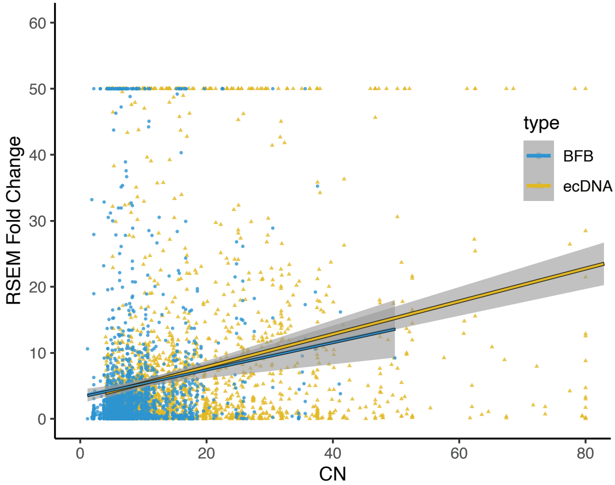

RSEM Fold Change
60
50
40
30
20
10
0
0
20
40
60
80
CN
type
BFB
ecDNA

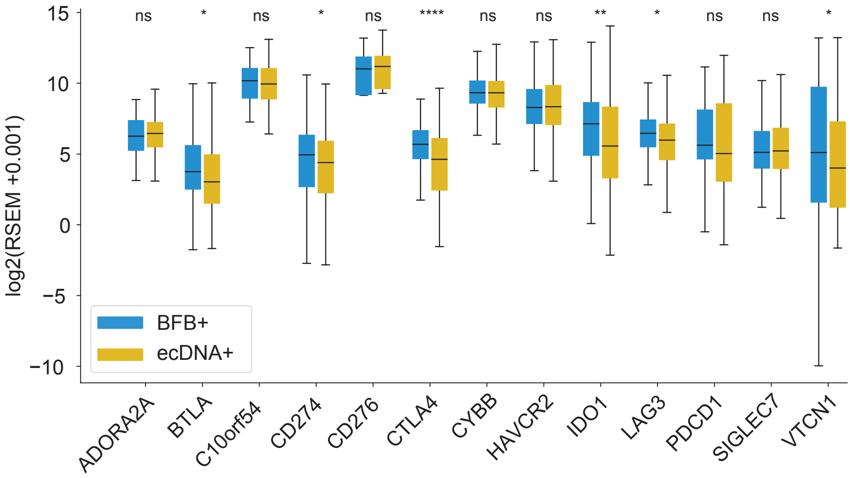

ns
*
ns
*
ns
****
ns
ns
**
*
ns
ns
*
15
10
5
0
−5
−10
log2(RSEM +0.001)
BFB+
ecDNA+
ADORA2A
BTLA
C10orf54
CD274
CD276
CTLA4
CYBB
HAVCR2
IDO1
LAG3
PDCD1
SIGLEC7
VTCN1

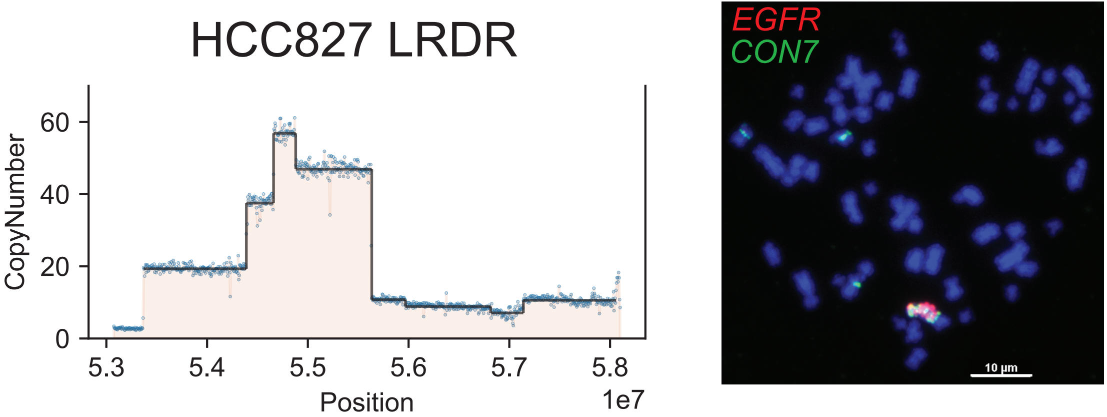
HCC827 LRDR
CopyNumber
0
20
40
60
5.3
5.4
5.5
5.6
5.7
5.8
Position
1e7
EGFR
CON7
10 µm

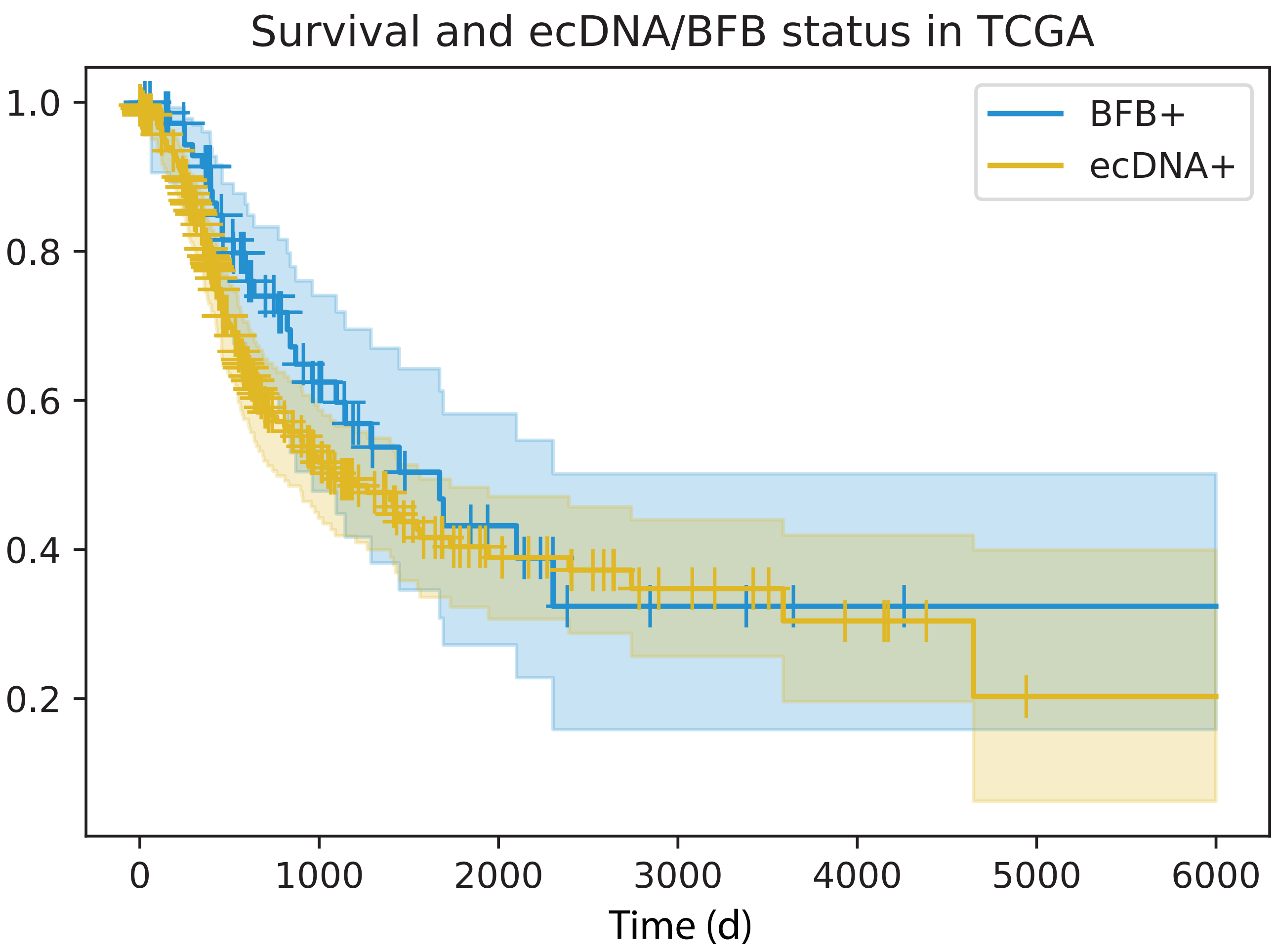
Survival and ecDNA/BFB status in TCGA
BFB+
ecDNA+
1.0
0.8
0.6
0.4
0.2
0
1000
2000
3000
4000
5000
6000
Time (d)

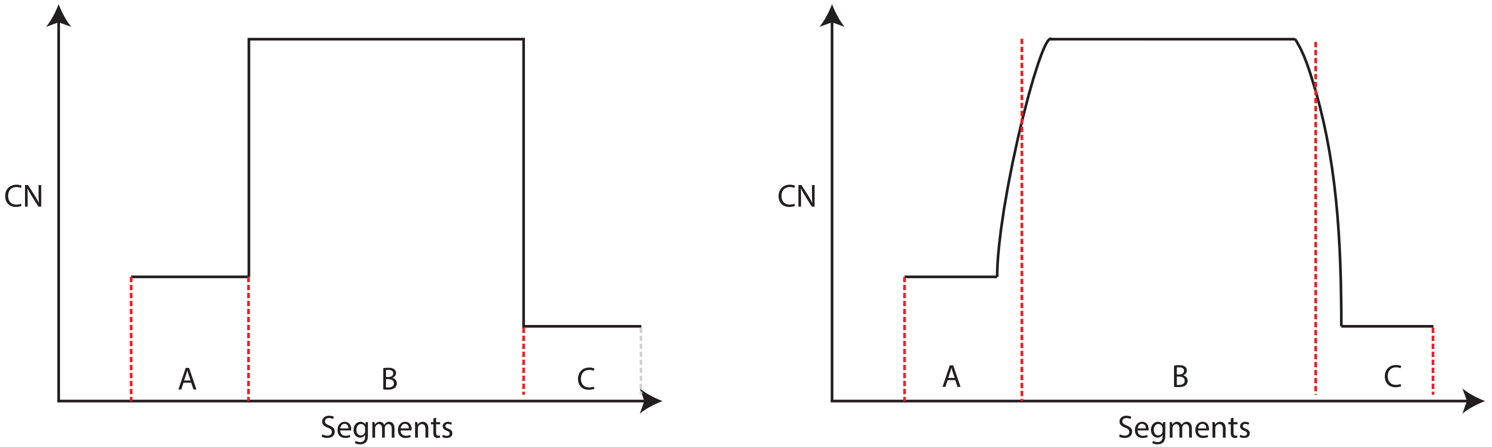
CN
A
B
C
Segments
CN
A
B
C
Segments
